# 50-year fire legacy regulates soil microbial carbon and nutrient cycling responses to new fire

**DOI:** 10.1101/2024.08.06.606581

**Authors:** Daniel Revillini, Christopher A. Searcy, Michelle E. Afkhami

**Author notes:** should be considered joint senior author (Christopher A. Searcy, Michelle E. Afkhami).

## Abstract

Fire disturbances are becoming more common, more intense, and further-reaching across the globe, with consequences for ecosystem functioning. Importantly, fire can have strong effects on the soil microbiome, including community and functional changes after fire, but surprisingly little is known regarding the role of soil fire legacy in shaping responses to recent fire. To address this gap, we conducted a manipulative field experiment administering fire across 32 soils with varying fire legacies, including combinations of 1-7 historic fires and 1-33 years since most recent fire. We analyzed soil metatranscriptomes, determining for the first time how fire and fire legacy interactively affect metabolically-active soil taxa, the microbial regulation of important carbon (C), nitrogen (N) and phosphorus (P) cycling, expression of carbohydrate-cycling enzyme pathways, and functional gene co-expression networks. Experimental fire strongly downregulated fungal activity while upregulating many bacterial and archaeal phyla. Further, fire decreased soil capacity for microbial C and N cycling and P transport, and drastically rewired functional gene co-expression. Perhaps most importantly, we highlight a novel role of soil fire legacy in regulation of microbial C, N, and P responses to recent fire. We observed a greater number of functional genes responsive to the interactive effects of fire and fire legacy than those affected solely by recent fire, indicating that many functional genes respond to fire only under certain fire legacy contexts. Therefore, without incorporating fire legacy of soils, studies will miss important ways that fire shapes microbial roles in ecosystem functioning. Finally, we showed that fire caused significant downregulation of carbon metabolism and nutrient cycling genes in microbiomes under abnormal soil fire histories, producing a novel warning for the future: human manipulation of fire legacies, either indirectly through global change-induced fire intensification or directly through fire suppression, can negatively impact soil microbiome functional responses to new fires.

## Introduction

As concerns surrounding the climate crisis and global carbon cycling continue to escalate in the Anthropocene, it will be vital to consider the impacts of major environmental disturbances on soil microbiomes, the biological drivers of many ecosystem functions (Delgado-Baquerizo et al., 2016, 2020; Falkowski et al., 2008). Fires are pervasive disturbances in terrestrial ecosystems and with environmental changes, shifting climatic patterns, historic fire suppression, and the accumulation of fuels, fires are becoming more unpredictable, more intense, and spreading further than in the past (He et al., 2019). While fire’s large societal impacts and strong effects on aboveground plant and animal communities are well established (Bowman et al., 2020; Certini et al., 2021), fire can have similarly important effects on soils, which hold nearly 80% of terrestrial carbon stocks and maintain >45% of global biodiversity (Anthony et al., 2023; Hawkins et al., 2023; Schimel, 2013). Additionally, soil microbes are the major engine of biogeochemical cycling on Earth; specifically, microbial functions underpin organic matter decomposition, carbon storage, nutrient cycling, and plant productivity (Bardgett & van der Putten, 2014). Fires can restructure soil microbial community composition and influence microbial functions with cascading effects on the direction and strength of these important ecosystem processes (Goberna et al., 2012; Hinojosa et al., 2019; A. R. Nelson et al., 2022; Yang et al., 2020), but the vast majority of current knowledge is based on observational, post-wildfire analysis of soils, which has obscured our ability to identify the *independent effects* (without confounding factors of time and environment) of fire on microbiome-mediated biogeochemical processes, including carbon (C) and nutrient cycling. Additionally, the majority of studies regarding fire effects on the microbiome have typically considered a range of weeks to years post-fire, but little is known about the immediate (i.e., within hours or days) responses in soils, which set the stage for post-fire successional dynamics to play out both below and aboveground. In a large-scale field inoculation study, Wubs et al., (2019) showed that the initial composition and functional capacity of the soil microbiome can shape the trajectory of plant community assembly for up to 20 years. While their study was not testing effects of fire, it highlights the long-term importance of early successional soil microbiomes on community assembly. Taken together, it is becoming apparent that to better understand the broad ecological effects of fire in the Anthropocene, we need to identify the direct and immediate impacts on soil microbial activity and functional capacity under realistic, yet controlled study conditions.

In addition, soil legacies – the collective history of abiotic and biotic influences on the soil (including soil microbial communities and functions) – can determine the ways soil, microbes, and associated plant communities respond to subsequent disturbances (Bakker et al., 2018; Kostenko et al., 2012; Scharer et al., 2023). Recent studies focusing on different global change factors have shown critical legacy effects from e.g., prior salinity and climate stress that shaped future plant-microbial interaction responses. For example, a legacy of high salinity for mangrove leaf- endophyte communities was essential for them to confer subsequent salinity-tolerance (e.g., increasing tree growth under high salt conditions), which increased mangrove performance (Subedi et al., 2022). Allsup et al., (2023) revealed that tree fitness increased when soil climate legacy and current climate stressors were aligned (e.g., microbiomes from warmer soils increased tolerance to current heat-stress in a suite of host trees). In a study focused on microbial responses, Canarini et al., (2021) found that recurrent droughts strongly shifted microbial communities and also enhanced soil multifunctionality, suggesting that soils with stronger drought legacies may increase their resilience to future droughts. Similar soil legacy effects may also occur for fire, especially in habitats that commonly experience fires or as fires continue to spread under global change.

Pyrogenic ecosystems have experienced many fire disturbances in the past; factors such as varying fire frequency and time since fire plausibly confer long-term effects belowground, altering soil structure, chemistry, resource availability, and the soil microbiome to varying degrees, yet we know very little about the ways fire and soil fire legacy interact to shape the soil microbiome. In the only study to our knowledge that has evaluated interactive effects of immediate fire and fire legacy, Revillini et al., (2021) analyzed fungal and bacterial compositional responses, but observed only strong pulse fire effects. Another study investigating the interaction between soil drought legacy and recent fire found that the recovery of burned soil microbial community composition and functional enzyme profiles to pre-fire levels was significantly slower for microbiomes with a drought legacy, with delayed recovery effects persisting for at least two years after disturbance (Hinojosa et al., 2019). Interestingly, methodology may play an important role in determining fire legacy effects, as studies have so far observed very few changes related to soil fire legacy using DNA amplicon sequencing or PLFA for microbiome characterization. To our knowledge, it remains unknown how fire legacy impacts microbial functional responses to new fires, and in order to identify these important and previously unreported interaction effects, targeted analyses of *active, functional* microbial responses are required.

Filling these gaps in our knowledge of how immediate and legacy effects of fire interact to shape microbiome function is crucial because of the numerous roles microbiomes have in ecosystem functions. Fires have pronounced impacts on biogeochemical cycling, including changes in pH, soil organic matter, nutrient availability, and nutrient leaching (Knicker, 2007; Pierson et al., 2019; Wan et al., 2001), which implies a strong concomitant effect on microbial functional responses after fire. Beyond the direct effects of heat from fire on soil microbial survival in upper soil layers, high-intensity fires can completely remove the typically consistent biomass inputs to soil from aboveground vegetation, increase inputs of pyrogenic forms of carbon, and also dramatically affect the forms and amounts of available nutrients through chemical transformations in the topsoil layers (Lasslop et al., 2019; Pellegrini et al., 2020, 2022). Taken together, these changes can greatly alter the functional niches available to the surviving, active soil microbiota. Previous studies have also observed significant increases in microbial genes involved in nitrogen (N) cycling after fire (e.g., *nif* – N2-fixation, *nar* – nitrate reduction, *nir* – nitrite reduction), presumably in response to the post-fire influx of various biomass-derived N compounds, which can be important for increasing plant germination and root development in N-limited systems (Dove et al., 2022; Hinojosa et al., 2019; Martin et al., 2022). It is important to note that all of these functional responses were measured on scales of weeks to decades after fire, and over time, environmental factors such as rainfall (leading to leaching) as well as plant community and soil biochemical changes likely influence microbial functional dynamics, making it difficult to specify independent effects of fire. Additionally, these transformations over time are particularly important in the context of variation in soil fire legacy, for example the effects of fire on soil C or N may compound with a greater number of historic fires or decline as time since fire increases (Li et al., 2021). Therefore, considering fire legacy interactions with recent fire, and a more immediate post-fire sampling of soils, will be helpful to provide greater clarity on the impacts of shifting fire regimes on soil microbial functioning, particularly for soil C cycling and sequestration (Pellegrini et al., 2022; Pellegrini & Jackson, 2020).

Soils, in fact, harbor the largest pool of carbon worldwide, with double the global biotic C pool and three times the global atmospheric C pool (Crowther et al., 2019). Fire has been shown to decrease the important soil C pool by ∼15%, as well as changing the types of carbon inputs into soil and shifting both soil respiration and microbial C degradation functions (Knicker, 2007; Li et al., 2021; Wang et al., 2012). These chemical transformations of C after fire can lead to a shift in availability of C forms (e.g., typically from labile to recalcitrant), with the potential to alter soil microbes, and ultimately impact important ecosystem functions they provide. As soil C resources change over time and after fire, presumably so will the functional niches for microbial taxa, filtering for particular C-cycling functions that may not be expressed in unburned, long-since burned, or infrequently burned soils (Pingree & DeLuca, 2017). An important feature of fire- deposited organic matter from the perspective of soil carbon storage is its presumably high chemical recalcitrance -- the ability to resist microbial or enzymatic degradation. Post-fire carbon forms are typically high C and low N, consisting of aromatic C rings and aliphatic C chains that are proposed to represent net ecosystem C sequestration, but recently Nelson et al., (2022) found evidence that soil microbial capacity to degrade pyrogenic aromatic carbons (catechol and protocatechuate) increased significantly after high severity forest fires. Catechol and protocatechuate were presumed to be highly recalcitrant forms of soil carbon persisting for decades to centuries (Pingree & DeLuca, 2017; Singh et al., 2012), and yet post-fire microbial interactions can have important and previously unpredicted effects on soil carbon storage (see Nelson et al., 2022). With continued human interventions, ever-increasing atmospheric CO2, warmer and drier conditions in many parts of the globe and more unpredictable fire regimes, it is more important than ever to understand microbial responses to fire, including the ways varying forms of post-fire C are processed, utilized, or stored.

In this study, we conducted a field experiment using prescribed fire and a unique set of 32 soils with a known fire legacy record dating back more than 50 years, and performed metatranscriptomic analyses to identify the immediate impact of high-intensity fire as well as the fire interaction with two important fire legacy metrics on metabolically active soil microbial communities and the expression of important belowground carbon and nutrient cycling functions. In unburned soils and those that experienced fire, we identified the immediate responses of metabolically active microbial taxa, individual microbial functional genes, carbohydrate-active pathways, as well as gene co-expression networks. Importantly, we were able to incorporate two aspects of soil fire legacy, time since last fire (TSF) and number of historic fires, to further determine how the microbiome’s past influence from fire (soil fire legacy) shapes its response to new fires, especially for critical soil microbial processes involved in carbon and nutrient cycling.

This experimental design allows us to identify microbial functional responses to recent fire that may be entirely dependent on a particular aspect of fire legacy or the combination of multiple aspects of fire legacy (i.e., interactions between recent fire, number of past fires, and time since the most recent fire). Additionally, the diverse combinations of fire legacy metrics and the interaction with recent fire, allow us to identify the effects of atypical soil fire legacies (e.g., recent fire suppression: many historic fires but long TSF) on microbiome responses to recent fire. A better understanding of the effects of these abnormal fire legacies is important as they are largely driven by anthropogenic global change or direct human interventions such as fire suppression (Jones et al., 2022; Santín & Doerr, 2016). Thus, we aimed to determine if (1) there is a strong, immediate effect of fire on soil microbiome functionality, and (2) how past fire history regulates microbial functional response to subsequent fires. Specifically, our goal was to evaluate the potential of fire and its interaction with fire legacy to disrupt belowground microbial C processing or to alter the cycling of growth-limiting nutrients. We hypothesized that *immediately after fire*, the soil metatranscriptome will reveal 1) shifts in which taxa are metabolically active, with taxa differentially up/down-regulated according to their functional niches and greater post-fire expression of fungal than bacterial genes, and 2) changes in expression of important soil C and nutrient cycling functional genes after the deposition of post-fire C compounds and nutrients, selecting for increases in targeted functions such as aromatic C degradation or nitrate reduction as well as disruption to metabolic pathways involved in glycolysis/gluconeogenesis or amino acid and citrate (TCA) cycling. Importantly, we also hypothesize that 3) *the interaction of soil fire legacy with recent fire* will strongly influence the expression of active microbial taxa and important C and nutrient cycling functions. The results will help us understand the potential of the post-fire microbiome to alter assembly dynamics, both below- and above-ground, through functional transformations of the resource environment for primary successional species including pyrophilic fungi or biocrusts, as well as pioneer annual plants. Given the concerning effects of both global change and human interventions on fire regimes, a greater understanding of the potential regulating role of soil fire legacy metrics for soil microbial functioning after fire is needed to form a clearer picture of fire ecology in the Anthropocene.

## Materials & Methods

### Site and study design

We conducted this study with soil microbiomes collected from the pyrogenic and imperiled Florida rosemary scrub habitat, which is dominated by patchily-distributed allelopathic rosemary shrubs (*Ceratiola ericoides*) interspersed with open-sand ‘gaps’ (Menges et al., 2008, 2018). Many rare and endemic plants depend on these gaps, which are maintained by fire (Menges et al., 2017). This habitat has a mean fire return interval of 20-59 years (David et al., 2019). Additionally, it was recently determined that the soil microbiome in this habitat is affected by pulse fire disturbance such as prescribed burning, and further that the post-fire soil microbiome reduces native-plant growth compared to unburned soil microbiomes (Revillini et al., 2021). Hernandez et al., (2021) found that environmental stress in this habitat led to declines in both bacterial and fungal diversity, and that increasing stress produced increasingly unstable microbial networks that may be more susceptible to disturbances, like fire. However, it remains unclear to what extent the function (rather than composition) of microbiomes is shaped by fire and the legacy of past fires.

### Soil collection and fire treatment

To understand microbial functional responses to fire and their fire legacy, we collected soils from 36 open sand gaps at Archbold Biological Station (ABS; 27°13392 N, 81°35051 W; Supplemental Table 1) with unique fire legacies that vary in time since fire (TSF) and historic number of fires. This detailed examination of fire legacies was enabled by fire records for ABS that began in 1967. Soil fire histories ranged in TSF from 1-33 years and historic number of fires from 1-7. Approximately 2.35 L of soil was collected from each of the 36 unique rosemary open sand gaps and placed in four foil trays (24 cm x 13 cm x 6.25 cm) using sterile technique and covered with aluminum foil. Importantly, we maintained soil structure during collection, with top soil layers from the field also at the top of the soil in trays. Three soil trays from each gap were transferred to a single rosemary scrub site at ABS to undergo a prescribed burn, and were separated into three replicates (burn blocks) to ensure uniform fire coverage across each block. One tray from each site was placed outside of the burn area as the ‘control’ treatment soil. To determine fire coverage and intensity, we placed nine HOBO temperature data loggers randomly across each burn block (Supplementary Figure 1). The prescribed burn was carried out on May 8, 2019. Within 4 hours of prescribed fire, 50 mL of soil was collected from the top 5 cm of each tray, flash frozen in the field with liquid N, and then stored at -80℃ until RNA extractions were performed.

### Soil metatranscriptomes

Total RNA was extracted from samples collected from the control samples and from the burn block that experienced the highest and most consistent temperatures (Supplemental Figure 1; n = 72) using the RNeasy PowerSoil Total RNA Kit (Qiagen, Carlsbad, CA, USA) and a slightly modified protocol. Briefly, we used ∼4 g of homogenized soil, 2.25 mL PowerBead Solution, and 3.25 mL phenol/chloroform/isoamyl alcohol (25:24:1) for extractions. Wash and elution steps were performed in an Avanti JXN-26 High-Speed Centrifuge (Beckman Coulter Inc., Brea, CA, USA) at 4500 g, and final elution was performed with 75 µL elution buffer. RNA was quantified with a Qubit 4 fluorometer (Qiagen, Carlsbad, CA, USA) using the Qubit RNA HS Assay Kit. Forty-eight samples that met the sequencing facility standards were then sent to the Joint Genome Institute (Walnut Creek, California, USA) for further quality control and sequencing on Illumina NovaSeq 6000 (S4 PE). Libraries were prepared using Qiagen FastSelect using a minimum of 100 ng RNA after ribo-depletion.

### Bioinformatics

Fastq files were first de-interleaved using the reformat.sh command from BBmap (v39.0). Following the SAMSA2 workflow (Westreich et al., 2018), all de-interleaved fastq files were cleaned to remove low-quality sequences and adaptor contamination using the program Trimmomatic (v0.36), where phred = 33, sliding window = 4:15, and minlen = 70. This process retained ∼98% of all sequence reads per sample. SortMeRNA (v2.1) was then used to retain only mRNA for annotation, and DIAMOND (v0.8.38) was used to match mRNA sequences against the NCBI-nr database (accessed May 19, 2022) with blastx (-k 25). MEGAN6 Ultimate Edition (v6.24.16; Huson et al., 2016) was then used to aggregate all annotated reads (>154 million) to the RefSeq database for taxonomy, and the KEGG database for functional genes (KEGG orthologs, genes) from the MEGAN6 UE database mapping file (<u>February 2022</u>). Additionally, to identify all transcripts associated with carbohydrate-active enzymes (CAZymes), mRNA sequences were matched against the CAZy database (accessed August 06, 2022) using DIAMOND. MEGAN6 was then used to identify matching taxonomy and functional genes (genes) associated with the CAZy database annotated reads (>1.8 million). All prior steps were performed using the Northern Arizona University <u>Monsoon compute cluster</u>. Finally, read counts were extracted from MEGAN6 for taxa at the phyla level, and for all known functional genes at the KEGG orthologs (KO; functional gene identifier) level for further analyses described below.

### Data Analysis

#### Metatranscriptome-wide taxonomic and functional gene expression patterns

To evaluate the broad effects of immediate fire and soil fire legacy (time since fire (TSF) and historic number of fires) on active microbial taxa we performed a distance-based redundancy analysis (db-RDA) with Bray-Curtis dissimilarity on the expression activity of all microbial phyla using the individual and interaction terms of three factors (fire treatment, TSF, and number of fires) as explanatory variables. Another db-RDA with the same explanatory variables but with expression across all functional genes as a response variable was used to investigate broad-scale changes in microbiome functional expression resulting from fire, fire legacy (TSF and number of fire), and their interactions. For the functional genes analysis, the original 4,552 KO set was first filtered to remove all samples with low expression variance (<55% of the mean; see Lemoine et al 2023) using the *filter_low_variance* function in the GWENA package in R (Lemoine et al., 2021; R Core Team, 2020) resulting in a total of 2,581 functional genes. Bray-Curtis dissimilarity matrices were calculated using the *avgdist* function from the vegan package. For the db-RDA of all active taxa and all functional genes, samples were rarefied to maintain those with >296,000 taxa reads, or >1,500 functional gene reads, respectively, resulting in 33 total samples for each analysis (18 unburned soils and 15 burned soils). Finally, to compare the amount of variation in active microbial communities between the burned and unburned soils, multivariate homogeneity of group dispersion from mean centroids was evaluated using the *betadisper* function.

#### Microbial community responses to recent fire and soil fire legacy

After detecting community-wide changes in which microbial phyla showed metabolic activity in response to immediate fire and fire legacy, we investigated which specific microbial groups responded to the experimental treatments by performing linear mixed models (LMM) on the centered log-ratio (CLR)-transformed phylum abundances for the 64 microbial phyla found in our samples using the *lmer* function from the lme4 package. With this LMM, we determined the main and interactive effects of recent fire, TSF, and historic number of fires, and included a random effect of soil collection site (gap = the specific open sand habitat patch): *CLR*_taxa_∼*burning*×*TSF*×*num*.*fires* + (1|𝑔𝑎𝑝). After correcting for multiple comparisons using the Benjamini- Hochberg procedure, we identified 31 metabolically active microbial phyla that responded to experimental fire, soil fire legacy metrics, and/or their interaction.

#### Critical carbon and nutrient cycling gene expression responses to fire and soil fire legacy

To determine how microbial functional genes respond (up- or down-regulation) to recent fire disturbance as well as the two important fire legacy metrics (TSF and num. fires), and their interactions, we used a for loop that performed LMMs on CLR-transformed functional gene abundances of targeted carbon (C), nitrogen (N), and phosphorus (P)-cycling KEGG genes. For our targeted analysis of functional genes, we subset the abundances of 124 important C-, N-, and P-cycling functional genes associated with critical biogeochemical processes (e.g., C degradation, C metabolism, N-fixation, or nutrient acquisition and transfer) from the full set of 2,581 genes. The target C, N, and P genes were identified *a priori* from literature regarding soil microbial functional genes where either gene names or KEGG orthologs were present (Garcia-Pausas et al., 2022; Goberna et al., 2012; Ludwig et al., 2018; A. R. Nelson et al., 2022); see Supplemental Table 2). The model was the same as above, but with the *a priori* selected C, N, and P functional genes as the responses: 𝐶𝐿𝑅_CNP_ ∼ 𝑏𝑢𝑟𝑛𝑖𝑛𝑔 × 𝑇𝑆𝐹 × 𝑛𝑢𝑚. 𝑓𝑖𝑟𝑒𝑠 + (1|𝑔𝑎𝑝). Additionally, we performed linear mixed models with the same explanatory variables but with carbohydrate-active enzyme pathways (Enzyme Commission number; EC) identified from the CAZy database as the response variable to gain insight into the potential selective effects of fire and fire legacy on active carbohydrate-cycling enzymes expressed by the soil microbial metatranscriptome. All LMM results are presented after correcting for multiple comparisons using the Benjamini-Hochberg procedure.

#### Gene co-expression network responses to fire and fire legacy

To assess if fire and fire legacy caused rewiring of functional gene coexpression in the soil metatranscriptome, we performed weighted gene co-expression network analysis using the WGCNA and GWENA packages in R. This method allows us to: 1) determine genes whose expression changes in tandem using soil metatranscriptome-wide RNAseq data, 2) identify sets of co-expressed genes (modules) associated with particular experimental treatments of fire or fire legacy, and 3) reveal which functional pathways were enriched in the gene modules whose expression responded to fire and fire legacy. To do this, directed (‘signed’) co-expression networks were constructed using gene-gene Spearman’s correlation edge weights where R^2^ > 0.9. A complete network incorporating all genes from both unburned and burned soils was constructed, and modules were identified using a branch cut at 0.5 from the agglomerative hierarchical clustering tree (Langfelder & Horvath, 2008; Lemoine et al., 2021). Co-expression eigengenes (module eigengenes), which are the primary axis of variation in gene co-expression, were determined by the loadings of the first principal component of the module gene expression matrix, and the PC scores from this first principal component were then used to identify relationships (significant correlations) with fire and fire legacy treatments, thereby linking changes in co-expression to fire effects. Enriched KEGG metabolic pathways within the separate unburned or burned modules from the co-expression network were calculated using over-representation analysis (*enrichKEGG*, reference database set to ‘ko’) in the clusterProfiler package (v4.8.3; Yu et al., 2012) to determine which microbial functional pathways (and associated genes) were most strongly represented with and without experimental fire. Additionally, separate co-expression networks were constructed using samples from either unburned or burned soils (also constructed using R^2^ > 0.9 and module merges at 0.5), to determine the way that fire might increase or decrease the number of co-expressed modules, providing insight for the coordination of functional gene sets in soil. For example, fire could either increase modularity through addition of new functional niches or decrease modularity by halting microbial activity and limiting the co-expression of genes. All statistical analyses were performed in R (v4.3.0) unless noted otherwise.

## Results

### Metabolically active prokaryotes respond more positively to fire than eukaryotes

The db-RDA of the active microbial community including 50 archaeal, bacterial and fungal phyla (after variance-filtering) revealed a significant effect of prescribed fire shaping composition (Figure 1; F1,27 = 8.34, *P* = 0.002), and also a main effect of historic number fires (F1,27 = 2.97, *P* = 0.049). Importantly, fire not only shifted the composition of metabolically active taxa, but also selected for a more homogeneous subset of taxa in which dispersion was greatly reduced (*betadisper*-F1,32 = 7.54, *P* = 0.009). From the LMMs, we reveal that of the 17 metabolically active microbial taxa that responded significantly to experimental fire treatment in this study, three active fungal phyla – Basidiomycota, Mucoromycota, and Chytridiomycota – greatly decreased expression relative to the unburned, controls (Figure 3; -60%, -52%, and -27%, respectively), while we observed increased expression across all responsive bacterial and archaeal phyla with the exception of Verrucomicrobia, which decreased expression after fire by ∼10% (Figure 2, Supplemental Table 3). We identified 12 bacterial and two archaeal phyla (Crenarchaeota and Thaumarchaeota) that significantly increased activity within hours of fire, including the increased expression of Actinobacteria, Bacteroidetes, Firmicutes, Deinococcus Thermus, and eight uncultured *Candidatus* bacterial phyla. Of the prokaryotic phyla that increased expression after fire, *Candidatus* Peregrinibacteria, Crenarchaeota, and Thaumarchaeota had the strongest responses in our model, with increases in phylum-level gene expression of 78%, 55%, and 50%, respectively.

**Figure 1.**
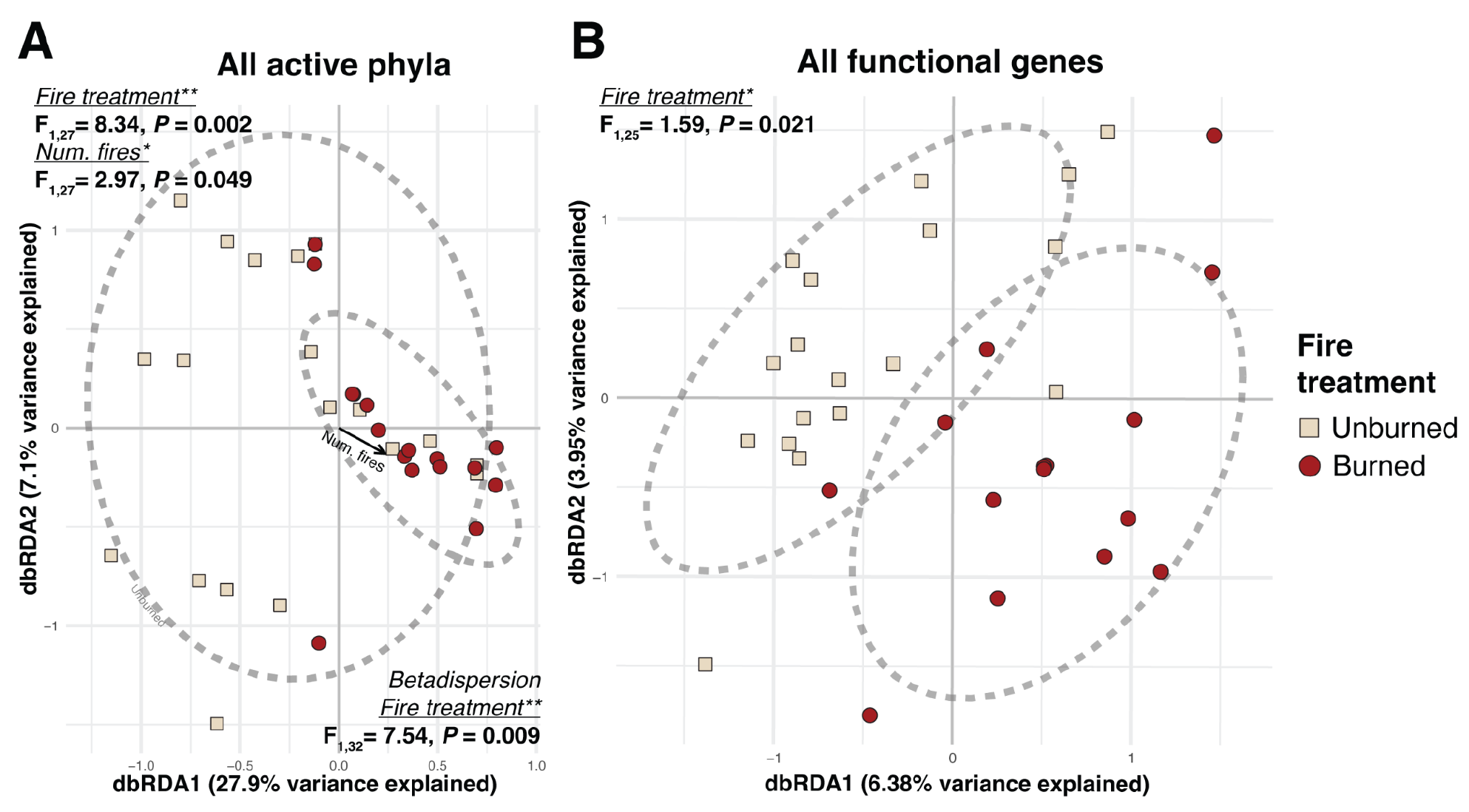
Fire effects on (A) metabolically active microbial taxa and (B) microbiome-wide functional gene expression. Bray-Curtis dissimilarity distance-based redundancy analysis for metabolically active archaeal, bacterial, and fungal phyla (n = 50, A), and variance-filtered functional genes (n = 2,581, B) are colored by fire experimental treatment, where soils were either unburned controls or experienced experimental fire. Fire not only shifted metabolically active taxa composition, but also selected for a more homogenous subset of taxa (significantly lower dispersion). Dashed lines represent 95% confidence intervals around fire treatment centroids.

**Figure 2.**
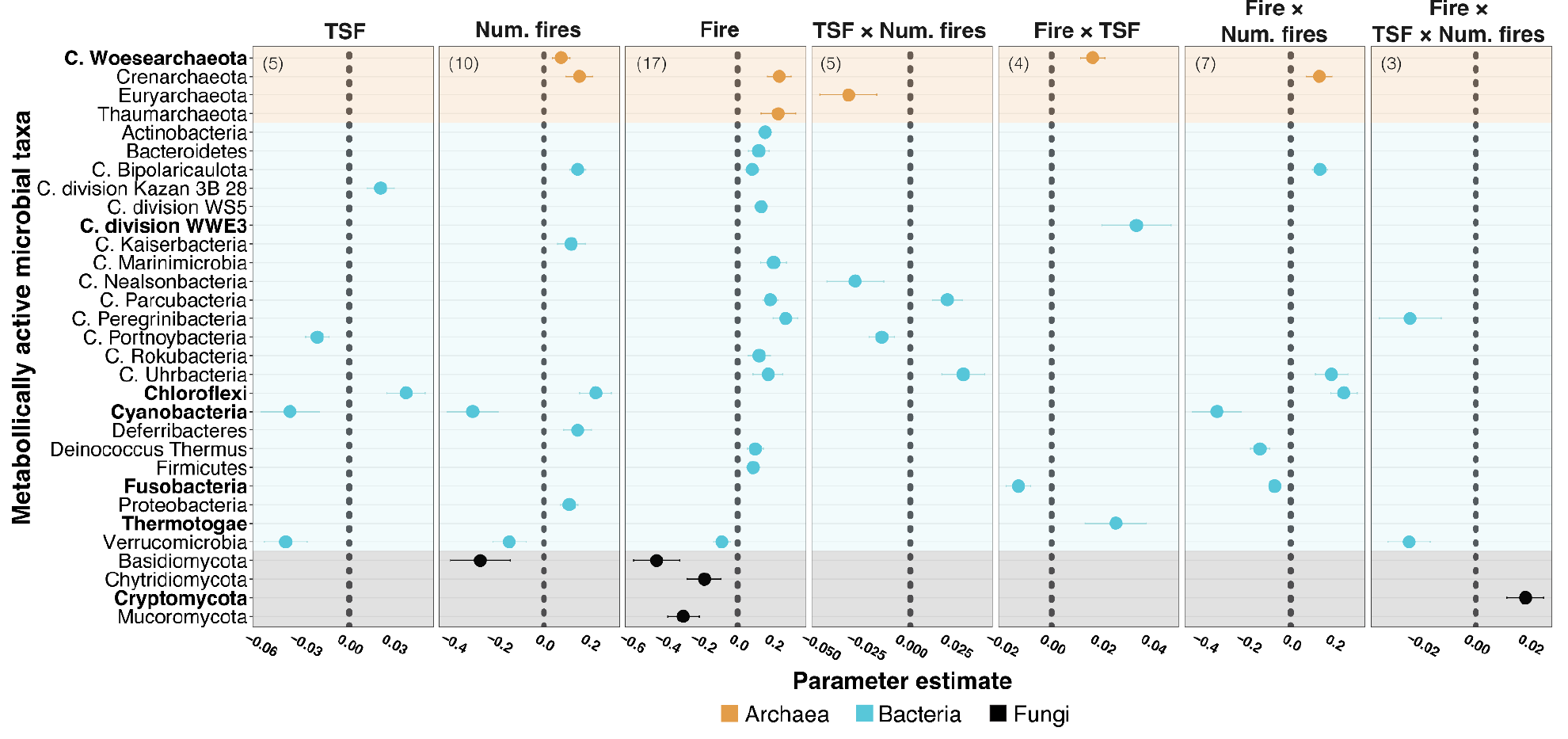
All metabolically active microbial phyla that responded significantly (P < 0.05 after correction for multiple comparisons) to recent fire, fire legacy, and their interactions. Plotted parameter estimates are from linear mixed models testing the response of CLR-transformed metabolically active microbial phylum abundances. Fire legacy metrics include time since fire (TSF) and number of historic fires (Num. fires). Response to the prescribed fire treatment (Fire) and its interactions with both aspects of fire legacy are plotted as well. Taxa with a ‘C.’ indicate *Candidatus* status. Taxa are colored by domain, total significant response counts are listed in parentheses (top left of each panel), and phyla that only responded to recent fire when considering soil fire legacy interactions are in bold. Fungal taxa had a notably different (negative) response to recent fire compared to the positive response of Bacteria and Archaea, and we observed that activity of ∼42% of all responsive taxa were significantly affected by fire x fire legacy interactions, highlighting an important effect of soil fire legacy on active microbial taxa after fire.

**Figure 3.**
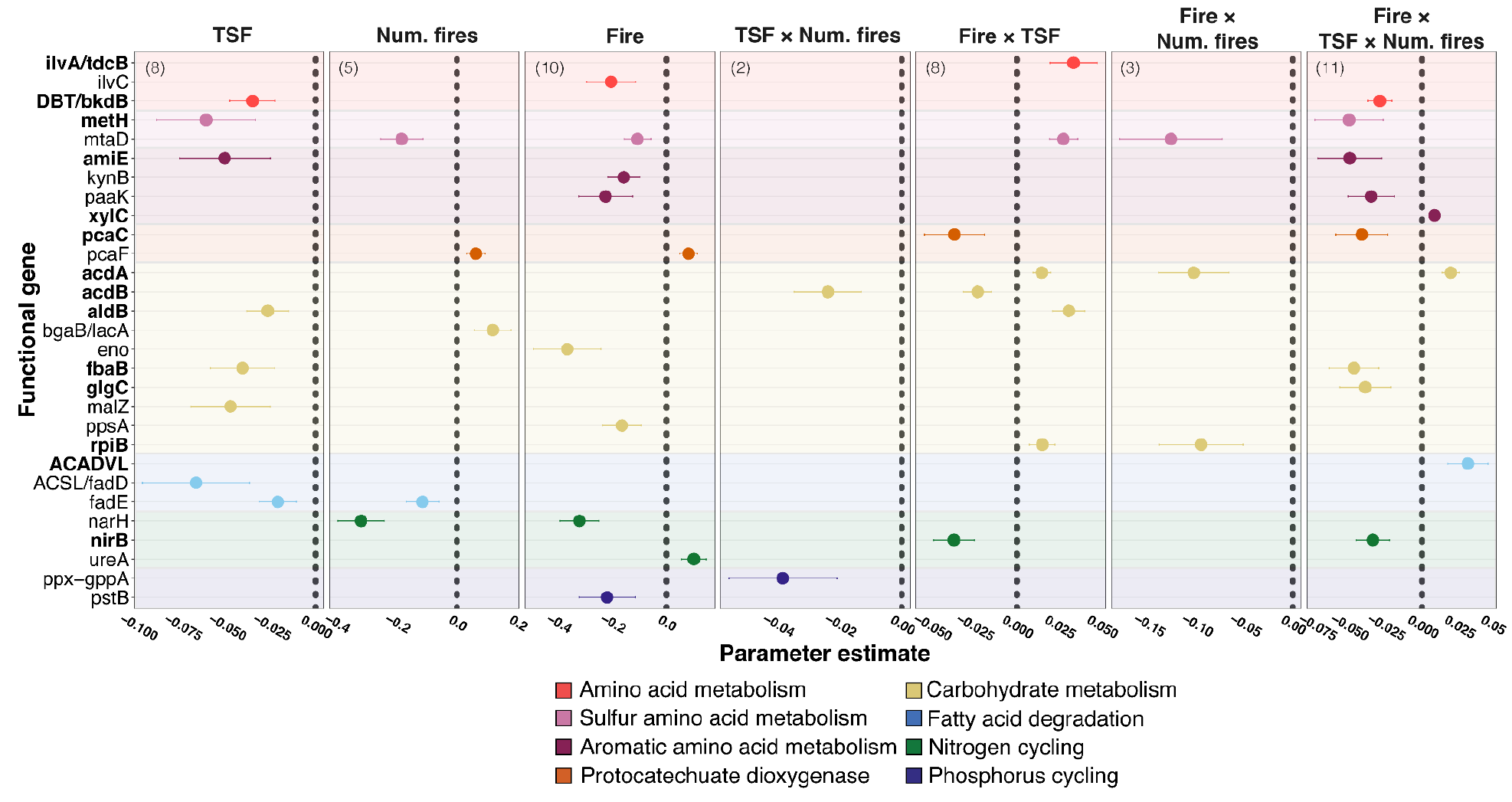
All critical carbon-, nitrogen- and phosphorus-cycling genes that responded significantly to recent fire, fire legacy, and their interactions. Plotted parameter estimates are from linear mixed models (LMM) testing the response of individual CLR-transformed KEGG ortholog abundance (functional genes); points represent significant gene expression responses to fire treatment, fire legacy, or their interactions (P < 0.05 after correcting for multiple comparisons). Fire legacy metrics include time since fire (TSF) and number of historic fires (Num. fires). Response to prescribed fire (Fire) and its interaction with fire legacy metrics is plotted as well. Genes are colored by higher-level functional groupings, and total significant response counts are listed in parentheses (top left of each panel). Functional genes exhibited primarily negative responses to recent fire and the three-way interaction term, which suggests strong negative effects of fire in the context of an atypical fire legacy (e.g., many historic fires, but long TSF). We show that the expression of 55% of these functional genes responded to fire x fire legacy interactions, and importantly 14 of 29 total genes (48%) *only* responded to recent fire in particular fire legacy contexts (genes highlighted in bold).

Soil fire legacy metrics had many main effects on the expression of active microbial taxa, and importantly, fire legacy (TSF and number of fires) influenced the response of active soil microbial expression to recent fire (see experimental fire interaction terms in Figure 2). These effects were more pronounced for prokaryotes than eukaryotes. In fact, similar to the effect of recent fire on prokaryotes, an increasing number of historic fires resulted in positive expression from 77% of the metabolically active prokaryotic phyla. Across metabolically active taxa, we found 14 fire ✕ fire legacy interaction responses, with ∼93% of these from archaea and bacteria. Taxa including the Chloroflexi and Cyanobacteria had some of the strongest responses to soil fire legacy and recent fire interactions. In soils with many historic fires that also experienced recent fire we observed a ∼200% decrease in abundance of Cyanobacteria, and again there was no separate main effect of fire treatment. For active fungi, Cryptomycota expression increased *only* under the three-way interaction of recent fire ✕ TSF ✕ num. fires (Figure 2), which indicates that the expression of Cryptomycota, a deeply divergent fungal phyla, is positively affected by new fire in soil with an atypical soil fire legacy. These results indicate that the effect of fire on these large soil microbial clades is governed by past fire legacy, and it is only in this context that their responsiveness to fire can even be detected.

### Main effects of recent fire and soil fire legacy downregulated the majority of responsive microbial C, N, and P cycling genes

The db-RDA across the entire (variance-filtered) gene expression dataset of ∼2,600 genes revealed a significant effect of prescribed fire shaping functional gene expression (Figure 1; F1,25= 1.59, *P* = 0.021). In order to understand these effects, we considered the microbial genes specifically implicated in critical C, N, or P cycling roles in the LMMs, and found strong effects of TSF, recent fire, and soil fire legacy effects on functional expression after recent fire, particularly for those soils with atypical fire legacies. This last is detected based on the three-way interaction, since this indicates a non-additive effect of recent fire when fire legacies are either simultaneously many historical fires but long time-since-fire (indicative of fire suppression) or few historical fires but recent time-since-fire (indicative of fire intensification).

After correcting for multiple comparisons, ∼25% of the 124 critical C, N, and P genes (32 of 124 genes) responded significantly to fire legacy metrics and/or recent fire (Figure 3). Notably, 80% of responses to the experimental fire treatment were negative, including downregulation of genes involved in N and P cycling as well as amino acid and carbohydrate metabolism. For genes with roles in nutrient mineralization and mobilization, which can be particularly important after disturbance events such as fire that lead to pulse nutrient releases in soil, we observed diverging expression patterns after the experimental fire treatment. For *ureA*, a primary urea-nitrogen hydrolysis gene, fire increased expression, but we also found that the expression of the phosphate transporter gene, *pstB*, and a nitrate-reducing gene, *narH*, both significantly decreased after fire. The aromatic amino acid metabolizing *paaK* gene (coding for phenylacetate-CoA ligase) was also downregulated in response to recent fire. Expression of the enolase coding gene, *eno*, had the strongest negative response to the prescribed fire treatment with a decrease in expression of ∼60% (Figure 3; Supplemental Table 3). Enolase is a critical enzyme in glycolysis, converting 2- phosphoglycerate to phosphoenolpyruvate, and paired with the decreased expression of *ppsA* (coding for phosphoenolpyruvate synthetase) after fire, it appears that fire may have an immediate impact on microbial capacity to produce ATP from glucose. In contrast, the experimental fire treatment increased *pcaF* gene expression by ∼50%, and this gene coded for beta-Ketoadipyl CoA thiolase, which suggests increased microbial aromatic carbon assimilation immediately after fire (Figure 3). Alternatively, *pcaC* (4-carboxymuconolactone decarboxylase), another gene that catalyzes the degradation of recalcitrant protocatechuate, was *only* responsive to fire in the context of fire legacy (Figure 3), further suggesting an important role of fire legacy in eliciting diverging responses from genes involved in similar functional roles in soil.

Soil fire legacy metrics generated varying responses from C, N, and P genes, but in general, gene expression was downregulated as fire legacy metrics increased for both TSF and number of fires. The main effect of TSF elicited the most gene responses of the two fire legacies in our study, and interestingly 100% of the responsive genes were downregulated with longer time since fires. These downregulated genes have previously been linked to either amino acid metabolism, carbohydrate metabolism, or fatty acid degradation (Figure 3), which indicates that as TSF increases, a wide range of important C-cycling genes will become less active. For example, two genes involved in different parts of the fatty acid degradation pathway (*ACSL/fadD* and *fadE*) are downregulated as TSF increases, as well as two genes involved in glycolysis (*fbaB* and *aldB*).

As the historic number of fires increased, we found that gene expression associated with amino acid metabolism and fatty acid degradation decreased as well as particularly strong downregulation of *narH* (59% decreased expression), an important nitrate-reducing gene (Figure 3). Only two genes responded positively to the historic number of fires: *pcaF,* which is associated with the degradation of protocatechuate, and *bgaB/lacA*, which facilitates the breakdown of low-molecular-weight carbohydrates to produce glucose. The ∼250% increased expression of *pcaF* in microbiomes with a legacy of many past fires suggests a higher capacity to metabolize aromatic carbon in soils that have experienced more historic fires (Chaleff et al 1974, Fischer et al 2021), while increases in *bgaB/lacA* expression could indicate that there is a higher turnover of soil C forms, including lactose, in microbiomes experiencing frequent fires.

### Fire legacy has a strong role in shaping microbial carbon, nitrogen, and phosphorus cycling functional responses to recent fire

Expression of 58% of the fire-responsive genes (or 14% of all genes examined) were affected by the interaction between our experimental fire and at least one aspect of fire legacy, suggesting microbial functional responses to new fire depend on the microbiomes’ fire legacy. Interestingly, the three-way interaction of experimental fire treatment with both aspects of fire legacy – TSF and historic number of fires – produced the greatest number of significant changes in functional gene regulation (38% of responsive genes and 9% of all genes examined, which is more response than any main effect of fire or any other interaction), and these genes represented almost all higher-level functional categories (Figure 3). In the LMMs, the three-way interaction term represents the microbial functional response to fire in the context of atypical soil fire legacies (i.e., infrequently burned soils that burned recently or frequently burned soils that have not been burned recently) that may occur more often under human influence (*i.e.,* fire intensification from climate change or disrupted fire regimes due to fire suppression, respectively). Expression of approximately 70% of these genes was downregulated in response to a new fire (Figure 3), including downregulation of genes implicated in glycolysis (*fbaB*), sucrose metabolism (*glgC*), aminobenzoate degradation (*amiE*), and degradation of pyrogenic protocatechuate (*pcaC*). This suggests significant downregulation of important functional genes after fire when soil microbiomes have an atypical combination of fire legacy. Additionally, all but one of these genes did not respond to the main effect of fire, suggesting again that microbial functional response to fire may be very context-dependent.

The interactive effects of fire and fire legacy on expression of critically important microbial functional genes reveal interesting shifts in the forms of C processed and impacts on microbial dissimilatory nitrate reduction in soils (Figure 3). For microbiomes that had a longer TSF legacy, recent fire significantly downregulated the nitrite reductase gene, *nirB*, by ∼89% (t = -3.1, *P* = 0.002). We also found that the expression of a gene responsible for metabolizing pyrogenic protocatechuate, *pcaC*, decreased significantly in soils with a longer TSF after recent fire (-62%; t = -2.07, *P* = 0.038). Two genes involved in glycolysis, *acdA* and *acdB*, responded differently to recent fire in soils with an increasing TSF, with *acdA* expression increasing by 37.5% and *acdB* expression decreasing by ∼130% (t = 2.87, *P* = 0.004; t = -2.75, *P* = 0.006, respectively). We identified upregulation of five amino acid and carbohydrate metabolism functional genes that are involved in the degradation or assimilation of more labile C forms under the interaction of recent fire and increasing TSF. These genes range in their functional roles in soil, from serine metabolism (*ilva/tdcB*) to glycolysis/gluconeogenesis (*acdA* and *aldB*), but interestingly, four of these five C-cycling genes *only* responded to recent fire in the context of increasing TSF.

### Downregulation of carbohydrate-active enzyme expression after recent fire and interactive effects of microbial fire legacy

The analysis of enzymatic expression pathways revealed that the main effect of fire leads to decreased expression of two important carbohydrate-cycling enzymatic pathways: carboxylic ester hydrolases and methyltransferases (Figure 4). These carbohydrate processing enzyme pathways are responsible for the degradation of hemicellulose and pectin in soil, respectively, and are mostly expressed by fungi and bacteria (Lombard et al. 2014). Methyltransferases also were interactively affected by the interaction of fire and the two soil legacy metrics (t = -2.46, P = 0.024), indicating that this carbohydrate-cycling enzyme pathway is further downregulated by fire when the microbiome has an atypical fire legacy. The responses of polysaccharide lyase and serine endopeptidase pathways to recent fire were likewise dependent on soil fire legacy (Figure 4 in bold). Polysaccharide lyase expression declined after recent fire *only* for microbiomes with a longer TSF legacy. Polysaccharide lyases are commonly associated with the fungal breakdown of lignin in soils (Kashewar 2017), therefore this result suggests that the ability of microbes to process lignin (one of the two chief constituents of wood) after fire is hampered in soils that have not experienced fire for many years. Expression of the serine endopeptidase pathway, which is associated with cellulolytic processes in soil and is often performed by bacteria (Yadav et al. 2019), *only* increased after fire in microbiomes that had experienced frequent fires in the recent past (Figure 4, in bold). Interestingly, some of the critical C, N, and P functional gene responses to the main effect of experimental fire can be related to the patterns observed for these carbohydrate- cycling enzymatic pathways, where, for example, declines in *eno* and *ppsA* genes (Figure 3) are associated with the decreased expression of the carboxylic ester hydrolase pathway, which all play important roles in glycolysis and gluconeogenesis.

**Figure 4.**
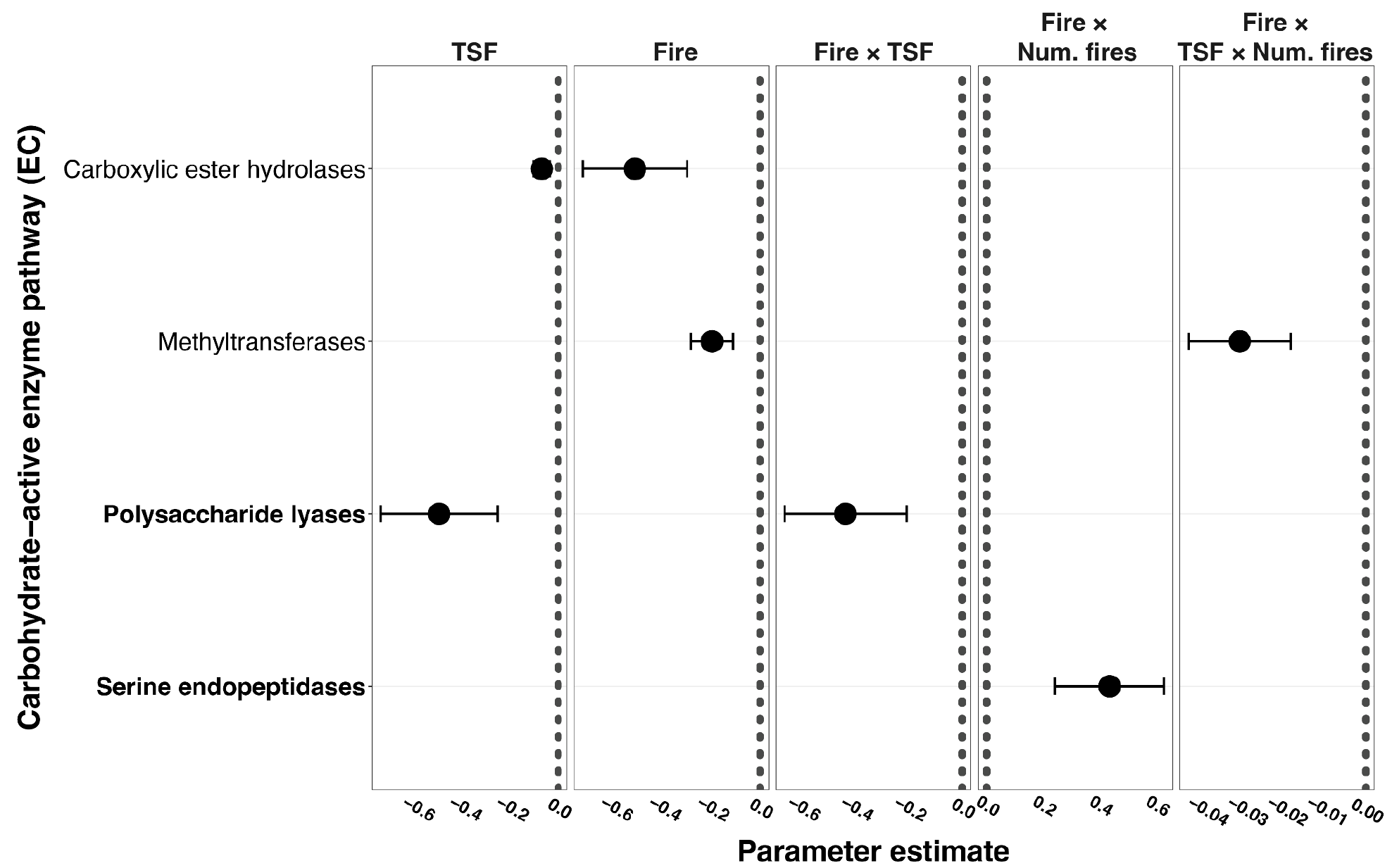
Carbohydrate-active enzyme (CAZyme) pathways that responded significantly (P < 0.05 after correcting for multiple comparisons) to fire,fire legacy, and/or their interactions. Plotted parameter estimates are from linear mixed models testing the response of CLR-transformed CAZyme pathways. Fire legacy metrics include time since fire (TSF) and number of historic fires (Num. fires). Response to prescribed fire (Fire) and its interactions with fire legacy metrics are plotted as well. CAZy pathways that only responded to recent fire in the context of certain fire legacies are in bold. CAZy pathways exhibited similar, negative responses to TSF and recent fire as observed in many critical carbon cycling genes, with implications for glycolysis and thermal stress responses.

### Microbiome functional gene co-expression networks were disrupted by fire

The microbial co-expression network of functional genes responsible for important metabolic pathways was significantly rewired by recent fire. We found that 44.6% of the 2,581 microbial genes in our network were in one of the three modules identified as rewired by fire. Specifically, we found that the largest module (#1), consisting of 816 functional genes, was upregulated in unburned soils, while module #5, consisting of 264 genes, was upregulated after experiencing the fire treatment. Hereafter, these modules are referred to as the unburned and burned modules, respectively. We also identified a functional gene module (#7) consisting of 132 genes that is positively correlated with increasing time since fire (Supplemental Figure 2). When we constructed co-expression networks of unburned and burned soils separately, we identified double the number of modules in the unburned network as in the burned network (i.e., 12 vs. 6 modules, respectively; Supplemental Figure 4). Collectively, these results indicate that soils exposed to recent fire maintain far less co-expression of functional genes (lower gene count in the burned compared to the unburned module), maintain fewer functional clusters than unburned soils (fewer modules in the burned network), and may have substantive disruption to some functional regulatory pathways (declined C-cycling pathway enrichment in the burned module).

We identified significantly enriched KEGG metabolic pathways within the two fire treatment modules (unburned and burned) to identify the potential rewiring of microbial functional co-expression networks after fire. Overall, 62 enriched KEGG pathways were identified in the unburned module compared to 25 enriched pathways in the burned module (Supplemental Figures 5-6), again indicating that fire simplified, or possibly disrupted, microbial functioning in the soil. In the unburned module, the carbon metabolism pathway was dominant among the 62 pathways (most representative within the module and composed of the most genes (n = 79; Figure 5). Additionally, we found that this module also maintained the greatest number of genes (545) and percentage of genes (52%) shared between functional pathways (i.e., genes contributing to multiple pathways). The substantially smaller burned module had 81 genes shared between functional pathways, representing 42% of genes. Together, these results show that fire led to degradation of the largest and most coordinated gene module, that fewer total modules were retained post-fire, and that there was declined enrichment of the most representative functional pathway (carbon metabolism), which suggests that fire broadly leads to a decline in gene co- ordination, and ultimately affects carbon cycling stability (Figure 5, Supplemental Figures 4-7, Supplemental Table 4).

**Figure 5.**
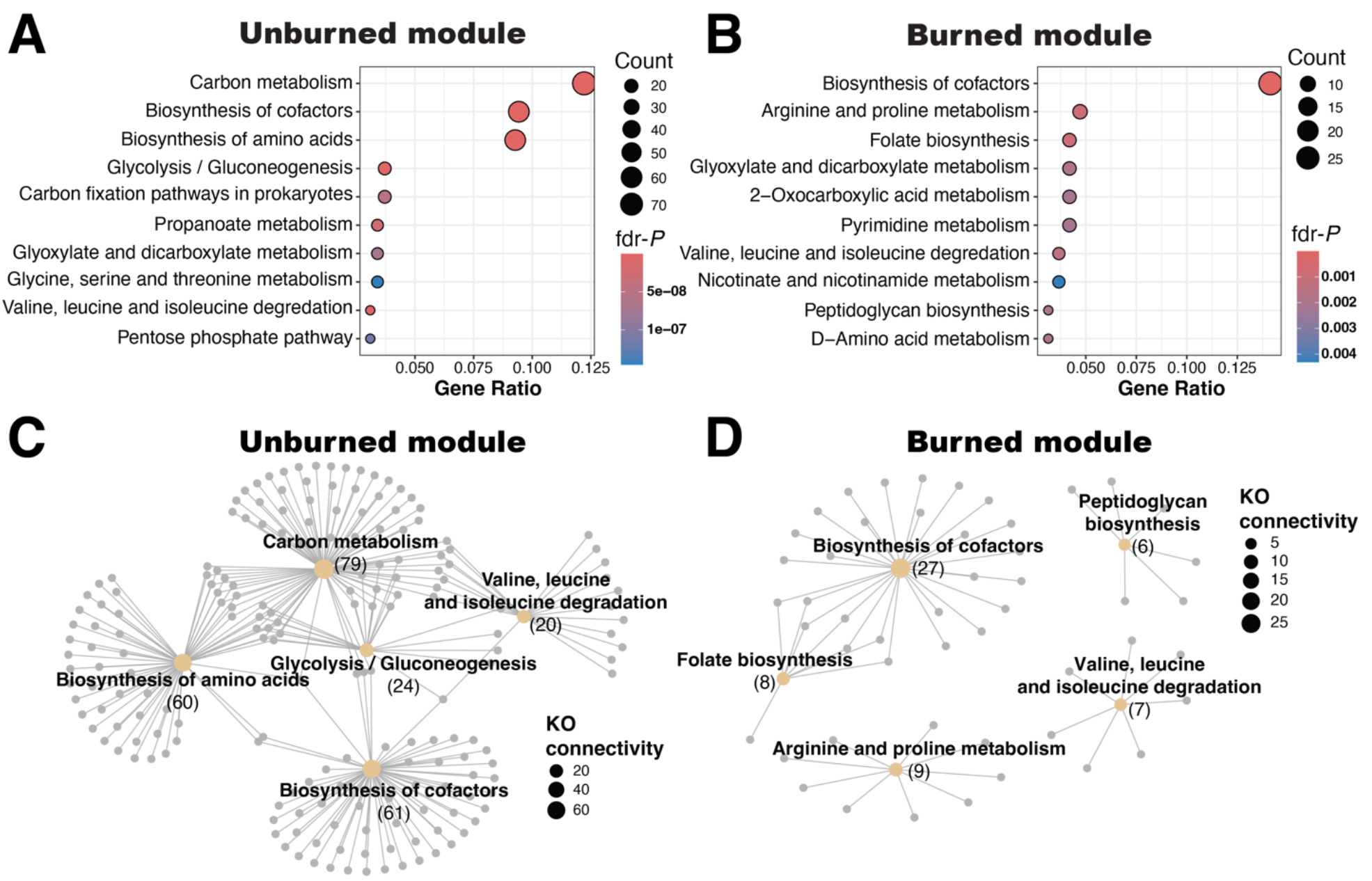
Functional pathways enriched in weighted gene co-expression network modules associated with unburned or experimentally burned soils. Gene ratios (observed pathway genes/total module genes) are plotted for the 10 most enriched pathways (lowest fdr-corrected P- connectivity of the top five enriched functional pathways in yellow and associated KEGG orthologs (genes) in grey for the unburned module (C) and the burned module (D). Note differences in count and P-value scales for A and B. The decrease in functional gene count and loss of many important carbon-cycling pathways (B), as well as the lack of functional pathway interconnectivity (D), indicate that fire strongly simplifies microbial functional co-expression networks and may destabilize carbon-cycling.

## Discussion

Fire disturbance had an immediate and strong effect on the soil metatranscriptome: shifting expression of metabolically active microbial taxa in the soil, reducing expression of carbohydrate- cycling enzymatic pathways, generally downregulating microbial C, N, and P cycling functions, and dramatically reshaping functional gene co-expression. Fire upregulated the expression activity of a wide range of prokaryotes immediately (13 of 14 fire-responsive prokaryotic phyla) while fungi showed the opposite response with downregulation of all three responsive fungal phyla (Figure 1). For gene co-expression networks, we found that fire decreased functional gene modules by 50% and resulted in decreased enrichment or complete loss of important coordinated C-cycling functional pathways (Figure 5; Supplemental Figures 4-7). We found clear evidence that soil fire legacy was critical in determining microbiome transcriptomic responses of active taxa, C, N, and P genes, and CAZy pathways to recent fire, where the interaction of fire and fire legacy metrics produced largely unique sets of taxa, functions, or enzymatic pathways from those observed when only considering a main effect of fire. Importantly, for soils with atypical fire legacies that then experienced recent fire, we observed both the greatest number of targeted C, N, and P gene responses, and that expression of these genes was almost entirely downregulated, which has implications for both future land management as well as microbiome functioning under a changing climate. Given these novel results, we emphasize the importance of keeping such legacy effects at the forefront in continuing efforts to fine-tune our understanding of microbiome responses to environmental disturbances like fire, and provide a caution to minimize further human-caused distortion of historic fire legacies.

### Microbiome responses to fire depend on soil fire legacy

Our results elucidate two major points regarding soil fire legacies: 1) that the human manipulation of fire legacies of soil (represented by our three-way interaction term) can dramatically impact the ways that the soil microbiome responds to recent fire and 2) the functional genes that responded significantly to recent fire × soil fire legacy interactions were almost completely different than those taxa and genes that responded to the main effect of fire (Figures 3). Towards the first of these points, the human influence on fire regimes takes different forms, and can be both indirect or direct (Kelly et al., 2020). Anthropogenic global change including land use change and climate change are clearly affecting historic fire regimes, generally leading to the intensification of fires (including their strength, coverage and frequency), but human interventions such as fire suppression or the increase in human-started wildfires are also altering fire legacies (Bowman et al., 2020; Jones et al., 2022). Few previous studies have attempted to understand the interactive role of fire legacy and recent fire in shaping biological responses, and to our knowledge, ours is the first study to investigate this effect on microbiome functioning. (J. E. D. Miller & Safford, 2020) analyzed the responses of plant communities to wildfires in habitats with a range of fire regime legacies and found that species richness recovery was greatest in regions where fire severity and return interval closely resembled the historic fire regime, suggesting that responses to recent fire were contingent on continuity with the fire legacy. Our results indicate that the expression of important geochemical cycling genes generally respond negatively in soils with atypical fire legacies that then experienced fire, suggesting that human alteration of fire regimes may limit the functional role of soil microbiomes.

Another striking outcome of our study was the strong role of fire legacy in regulating microbial functional responses to new fire. Given projected increases in environmental disturbances globally and their broad ecological impacts, it is important to measure direct responses to individual disturbance events such as drought, fire, or flooding, but given our results, it appears a crucial oversight to disregard the role of previous conditions (i.e., fire legacies) in shaping these responses. The concept of legacy effects interacting with current disturbance has been addressed in other systems, focusing on invasion ecology (A. D. Miller et al., 2021), land- use legacies (Bürgi et al., 2017), hydrological and precipitation legacies (Müller & Bahn, 2022; Tecon & Or, 2017), or summative work regarding legacy effects in forest ecosystem functioning (Perring et al., 2016), but few have pursued this question using an experimental system, and even fewer have investigated the role of microbiome function. The strong interactive effect of fire legacy and recent fire in our study was primarily observed for the critical C, N, and P functional genes rather than for which microbial taxa were active in the community. For these critical cycling genes, we identify 22 significant shifts for fire × fire legacy interactions, and only 10 in response to the main effect of fire. A striking finding was that 85% of the C and nutrient cycling genes that responded to fire × fire legacy interactions were unique from those that respond to recent fire (Figure 3), meaning that evaluating recent fire alone without considering historical context would miss not only the exact nature of the response, but would also fail altogether to identify many of the microbial functions that are altered by fire. Therefore, we propose that the incorporation of soil fire legacy effects into analysis of the immediate effects of fire is critical to holistically understand soil microbiome responses to fire in a changing world.

The way microbial functional expression responded to new fires in the context of soil fire legacy has important implications for the future of C and nutrient cycling under global change scenarios (Stephens et al., 2020). Altered fire regimes under climate change will create environments that depart in different degrees from their historical, and likely locally-adapted conditions (Jones et al., 2022; Naylor et al., 2020), presenting new scenarios that produce less-predictable community and functional responses. Given our results that show mostly negative microbial functional gene responses to recent fire, particularly in the context of soils that have atypical fire histories, we might expect these new fire regimes to also impact broader microbiome services such as C sequestration, decomposition, or N-cycling. As shifting fire regimes are paired with human interventions such as persistent fire suppression, it is more important than ever to determine how extensive microbial functional responses are, and whether they can impact ecosystem services provided by soils (Chen & Sinsabaugh, 2021; Qu et al., 2024; Yang et al., 2020). We particularly advocate for research that looks across ecosystems that range in natural fire dependency – from those that rarely experience fire to pyrogenic systems that frequently experience fire – and capture a gradient of abiotic variation in order to best understand local soil microbial adaptations in the context of fire legacy, and to better predict their responses to future disturbance.

### Archaeal and bacterial taxa increase expression after recent fire

The use of transcriptomics rather than amplicon sequencing in the current study highlighted some discrepancies in results of fire effects on active versus observed (active and relic) microbial communities in soil. In a previous analysis of the soils tested here, Revillini et al., (2021) found that bacterial rather than fungal community composition shifted immediately after fire, and that many bacterial taxa responded negatively to recent fire. In contrast, 13 of 14 responsive prokaryotic phyla increased their activity after fire when using a metatranscriptomics approach to evaluate changes in microbial functional gene expression. Additionally, we found that metabolically active Basidiomycota and Mucoromycotina decreased strongly after fire in this study (Figure 2). The negative and greater strength of fungal responses in this study (higher magnitude LMM estimates than from prokaryotes) align with many previous studies that predict stronger responses from soil fungi than bacteria to fire (Fox et al., 2022; Lucas-Borja et al., 2019; Pressler et al., 2019), though we did observe substantially more higher-level bacterial taxa that responded (almost all positively) to fire. Given these interesting positive responses of many bacteria and archaea to fire, it will be important to consider using methods including transcriptomics or qSIP that best determine the effects of pulse environmental disturbances like fire on active microbial taxonomic shifts and to reveal the clearest picture of fire effects belowground.

### Microbial carbon, nitrogen, and phosphorus functional responses to recent fire are broadly negative

Perhaps not surprisingly after a high-intensity fire, soil microbial functional gene expression generally decreased in our study, but these decreases were not all expected and suggest some important shifts in the way that microbes degrade carbon. The main effect of fire largely decreased the microbial genes that process C in soil, with downregulation of many amino acid and carbohydrate metabolizing genes, but led to the upregulation of the *pcaF* gene, responsible for the degradation of protocatechuate, a highly recalcitrant form of pyrogenic organic C. Interestingly, when considering the fire legacy of soils, we showed that the expression of many C-cycling genes also decreases as TSF increases. These findings suggest that not immediately after new fire but before a long TSF (33 years in our study) there might be an intermediate phase when soil microbes and associated functions are most active or efficient, which aligns well with previous studies investigating microbial enzyme activity and respiration responses to fire on short, medium and long-term (>30 years) time scales (Muñoz-Rojas et al., 2016; Pérez-Valera et al., 2019). Further, when soils with atypical fire legacies experienced new fire, expression for the majority of C genes decreased with the exception of *xylC*, *acdA*, and *ACADVL*, which are involved in catechol degradation, glycolysis/gluconeogenesis, and fatty acid transformation, respectively (Figure 3). This suggests that while the majority of microbial C-cycling will be negatively impacted, there is a range of C forms processed after fire in soils with an atypical legacy, from highly recalcitrant- to-labile. Ling et al., (2021) found that relatively shortly after wildfire (3 months), the boreal forest soil microbiome was largely processing recalcitrant C and leading to faster soil organic C (SOC) turnover, while 15 years after fire more labile forms of C were being processed. They hypothesized that the bacterial community becomes simplified after fire and these bacteria form a distinct relationship between the available pyrogenic OM (dominated by recalcitrant, aryl C) that shifts over decades to return to a more balanced system with more diverse SOC processing. Similarly, we observed a distinct homogenization and narrowing of the active microbiome in our study after fire, with significantly less community variation than found in unburned soils (Figure 1; *betadispersion*-*P* = 0.009), and also show that the expression of >90% of the bacterial taxa increased after fire, potentially pointing to an increased role for bacterial decomposition of pyrogenic OM after fires. Chemical recalcitrance of C is typically considered an important factor in calculating C storage potentials in soil, with higher proportions of recalcitrant C thought to correlate with greater soil C storage (Pingree & DeLuca 2017), but our results suggest that fire and soil fire legacy interactions can alter the ways that the soil microbiome can cycle both labile and recalcitrant C. This dynamic should be examined in greater detail to determine the effects of fire- fire legacy interactions on C sequestration and decomposition, important services largely regulated by the soil microbiome (Bahram et al., 2018; M. B. Nelson et al., 2016).

### Fire impacts the coordination of microbial functioning in soil

To identify broader scale implications for C, N, or P cycling, we used a gene co-expression network approach to better understand coordinated functional gene responses to fire. The co-expression network module for soils that experienced fire had fewer representative genes, less connectivity among functional pathways (see Supplemental Figure 7), and lost the significant enrichment of carbon-associated functional pathways that were dominant in the unburned control soils (Figure 5). Our results were similar to another study using experimental fire, but in grasslands, where they observed a dramatic decline in the relative abundance and connectivity of C degradation genes (Yang et al., 2020). In contrast, McClure et al., (2020) found that under drought stress the soil microbiome exhibited significantly increased community-wide functional coordination in the co-expression network. While drought and fire stress may impose similar pressures (heat, desiccation, etc.), fire disturbance is high intensity and short duration, whereas drought stress is typically recurrent, less dramatic and more prolonged, and this contrast between pulse and press disturbances may have a profoundly different effect on microbial functional coordination. In our study, the high-intensity fire treatment seriously disrupted connectivity of the microbial functional network, suggesting an overall lack of coordination among functional pathways, particularly for C-related functions. This breakdown in connectivity can lead to declines in C-cycling overall (Lopez et al., 2024), and limit the capacity to process a diversity of C forms in soil, which can be especially important after fire given the notable increases in C deposition and shifts in nutrient availability (Knicker, 2007; Pellegrini et al., 2020). The comparison of these pre- or post-fire co-expression networks revealed that the functional complementarity required to carry out important functions in soil such as glycolysis or the processing of complex aromatic C compounds may be lost after fire (Figure 5D; Supplemental Figure 7). One potential explanation for these functional breakdowns may be the fire-driven declines in fungal activity as suggested by Morriën et al. (2016), where the efficiency of carbon cycling can be related to shifts in the active fungal community within microbial networks. This finding suggests an interesting line of research to better understand the ways that microbial community shifts after fire can lead to changes in coordinated functional capacity that ultimately alters cycling of C and nutrients in soil for years into the future.

## Conclusion

Historic soil fire legacies can dictate the immediate taxonomic and functional microbial responses to fire, the beginning of a transformational process that carries on for decades-to-centuries (Dove et al., 2022; Pellegrini et al., 2022; Singh et al., 2012). Our results highlight 1) the major importance of a soil’s prior experience with fire in regulating current microbial responses to fire, and 2) that the vast majority of C, N or P-related functional gene responses to recent fire were negative, and particularly for soils with atypical fire legacy. The period we measured (within hours of disturbance) is critical for early colonizers and sets the stage for both below- and aboveground community assembly, but future work in this field would be well-served applying hourly-daily-weekly time series analyses. This would allow researchers to identify the functional transitions after fires over time and identify critical moments in post-fire succession related to decomposition, belowground diversity, and C and nutrient cycling. We identified multiple taxa and functions whose response to recent fire was *explicitly dependent on soil fire legacy*, suggesting that many important microbiome functional responses can be missed when researchers disregard the context of historic disturbance regimes, and we hope this insight influences the future research of environmental disturbances. Finally, we have shown that functional responses were particularly negative after fire for soils with atypical fire legacies. Thus, the impacts to soil ecosystem functions from fire may be most pronounced where fire has been suppressed for unnaturally long time periods or in ecosystems that had been previously unaffected by fire but where fire intensification is occurring, suggesting that land managers should pursue fire management strategies that most closely mimic historic regimes in order to minimize the loss of important microbiome functions. Globally, humans need to rapidly mitigate climate change to prevent further disruption of both fire regimes and adapted ecosystem property dynamics.

## Acknowledgements

Thanks to the fire management team at Archbold Biological Station, and especially to Kevin Main for coordinating prescribed burning in this project. Special thanks to Dr. Kasey Kiesewetter and Dr. Hunter Howell for their tireless support conducting field work. We are grateful for funding from the U.S. National Science Foundation and U.S. DOE Joint Genome Institute provided to Michelle Afkhami and Christopher Searcy, and to the Joint Genome Institute for providing metatranscriptomic sequencing service.

## Author Contribution Statement

**Daniel Revillini**: Conceptualization (equal); Investigation (lead); Writing – original draft (lead); Formal analysis (lead); Writing – review and editing (equal). **Christopher Searcy**: Funding acquisition (equal); Conceptualization (equal); Investigation (supporting); Formal analysis (supporting); Writing – review and editing (equal). **Michelle Afkhami**: Funding acquisition (equal); Conceptualization (equal); Writing – review and editing (equal).

## Data Availability Statement

All associated sequencing data uploads have been initiated in the NCBI Gene Expression Omnibus (GEO) database under accession: ASDFSDF. GEO upload includes all interleaved raw sequencing files from JGI as well as MEGAN6-processed KEGG ortholog counts per sample. Metadata, data analysis (R) script, and bioinformatics processing shell script have been uploaded to FigShare under project title: “*50-year fire legacy regulates soil microbial carbon and nutrient cycling responses to new fire*” with doi: 10.6084/m9.figshare.26203991.

## Artificial Intelligent Generated Content

There was no content generated using artificial intelligence in preparation of this manuscript.

## References

1. Allsup, C. M., George, I., & Lankau, R. A. (2023). Shifting microbial communities can enhance tree tolerance to changing climates. Science, 380(6647), 835–840. 10.1126/SCIENCE.ADF2027

2. Anthony, M. A., Bender, S. F., & van der Heijden, M. G. (2023). Enumerating soil biodiversity. Proceedings of the National Academy of Sciences, 120(33), e2304663120.

3. Bahram, M., Hildebrand, F., Forslund, S. K., Anderson, J. L., Soudzilovskaia, N. A., Bodegom, P. M., Bengtsson-Palme, J., Anslan, S., Coelho, L. P., Harend, H., Huerta-Cepas, J., Medema, M. H., Maltz, M. R., Mundra, S., Olsson, P. A., Pent, M., Põlme, S., Sunagawa, S., Ryberg, M., … Bork, P. (2018). Structure and function of the global topsoil microbiome. Nature, 560(7717), 233–237. 10.1038/s41586-018-0386-6

4. Bakker, P. A. H. M., Pieterse, C. M. J., de Jonge, R., & Berendsen, R. L. (2018). The Soil-Borne Legacy. In Cell (Vol. 172, Issue 6, pp. 1178–1180). Elsevier Inc. 10.1016/j.cell.2018.02.024

5. Bardgett, R. D., & van der Putten, W. H. (2014). Belowground biodiversity and ecosystem functioning. Nature, 515(7528), 505–511. 10.1038/nature13855

6. Bowman, D. M. J. S., Kolden, C. A., Abatzoglou, J. T., Johnston, F. H., van der Werf, G. R., & Flannigan, M. (2020). Vegetation fires in the Anthropocene. Nature Reviews Earth & Environment, 1(10), 500–515. 10.1038/s43017-020-0085-3

7. Bürgi, M., Östlund, L., & Mladenoff, D. J. (2017). Legacy Effects of Human Land Use: Ecosystems as Time-Lagged Systems. Ecosystems, 20(1), 94–103. 10.1007/s10021-016-0051-6

8. Canarini, A., Schmidt, H., Fuchslueger, L., Martin, V., Herbold, C. W., Zezula, D., Gündler, P., Hasibeder, R., Jecmenica, M., Bahn, M., & Richter, A. (2021). Ecological memory of recurrent drought modifies soil processes via changes in soil microbial community. Nature Communications, 12(1), 1–14. 10.1038/s41467-021-25675-4

9. Certini, G., Moya, D., Lucas-Borja, M. E., & Mastrolonardo, G. (2021). The impact of fire on soil-dwelling biota: A review. Forest Ecology and Management, 488(February). 10.1016/j.foreco.2021.118989

10. Chen, J., & Sinsabaugh, R. L. (2021). Linking microbial functional gene abundance and soil extracellular enzyme activity: Implications for soil carbon dynamics. Global Change Biology, 27(7), 1322–1325. 10.1111/gcb.15506

11. Crowther, T. W., van den Hoogen, J., Wan, J., Mayes, M. A., Keiser, A. D., Mo, L., Averill, C., & Maynard, D. S. (2019). The global soil community and its influence on biogeochemistry. In Science (Vol. 365, Issue 6455). American Association for the Advancement of Science. 10.1126/science.aav0550

12. David, A. S., Quintana-Ascencio, P. F., Menges, E. S., Thapa-Magar, K. B., Afkhami, M. E., & Searcy, C. A. (2019). Soil microbiomes underlie population persistence of an endangered plant species. American Naturalist, 194(4), 488–494. 10.1086/704684

13. Delgado-Baquerizo, M., Maestre, F. T., Reich, P. B., Jeffries, T. C., Gaitan, J. J., Encinar, D., Berdugo, M., Campbell, C. D., & Singh, B. K. (2016). Microbial diversity drives multifunctionality in terrestrial ecosystems. Nature Communications, 7, 1–8. 10.1038/ncomms10541

14. Delgado-Baquerizo, M., Reich, P. B., Trivedi, C., Eldridge, D. J., Abades, S., Alfaro, F. D., Bastida, F., Berhe, A. A., Cutler, N. A., Gallardo, A., García-Velázquez, L., Hart, S. C., Hayes, P. E., He, J. Z., Hseu, Z. Y., Hu, H. W., Kirchmair, M., Neuhauser, S., Pérez, C. A., … Singh, B. K. (2020). Multiple elements of soil biodiversity drive ecosystem functions across biomes. Nature Ecology and Evolution, 4(2), 210–220. 10.1038/s41559-019-1084-y

15. Dove, N. C., Taş, N., & Hart, S. C. (2022). Ecological and genomic responses of soil microbiomes to high-severity wildfire: linking community assembly to functional potential. ISME Journal, 16(7), 1853–1863. 10.1038/s41396-022-01232-9

16. Falkowski, P. G., Fenchel, T., & Delong, E. F. (2008). The microbial engines that drive earth’s biogeochemical cycles. Science, 320(5879), 1034–1039. 10.1126/science.1153213

17. Fox, S., Sikes, B. A., Brown, S. P., Cripps, C. L., Glassman, S. I., Hughes, K., Semenova-Nelsen, T., & Jumpponen, A. (2022). Fire as a driver of fungal diversity — A synthesis of current knowledge. In Mycologia (Vol. 114, Issue 2, pp. 215–241). Taylor and Francis Ltd. 10.1080/00275514.2021.2024422

18. Garcia-Pausas, J., Romanyà, J., & Casals, P. (2022). Post-fire recovery of soil microbial functions is promoted by plant growth. European Journal of Soil Science, 73(4), 1–15. 10.1111/ejss.13290

19. Goberna, M., García, C., Insam, H., Hernández, M. T., & Verdú, M. (2012). Burning Fire-Prone Mediterranean Shrublands: Immediate Changes in Soil Microbial Community Structure and Ecosystem Functions. Microbial Ecology, 64(1), 242–255. 10.1007/s00248-011-9995-4

20. Hawkins, H. J., Cargill, R. I. M., Van Nuland, M. E., Hagen, S. C., Field, K. J., Sheldrake, M., Soudzilovskaia, N. A., & Kiers, E. T. (2023). Mycorrhizal mycelium as a global carbon pool. In Current Biology (Vol. 33, Issue 11, pp. R560–R573). Cell Press. 10.1016/j.cub.2023.02.027

21. Hernandez, D. J., David, A. S., Menges, E. S., Searcy, C. A., & Afkhami, M. E. (2021). Environmental stress destabilizes microbial networks. ISME Journal, *1*. 10.1038/s41396-020-00882-x

22. Hinojosa, M. B., Laudicina, V. A., Parra, A., Albert-Belda, E., & Moreno, J. M. (2019). Drought and its legacy modulate the post-fire recovery of soil functionality and microbial community structure in a Mediterranean shrubland. Global Change Biology, 25(4), 1409– 1427. 10.1111/gcb.14575

23. Huson, D. H., Beier, S., Flade, I., Górska, A., El-Hadidi, M., Mitra, S., Ruscheweyh, H. J., & Tappu, R. (2016). MEGAN Community Edition - Interactive Exploration and Analysis of Large-Scale Microbiome Sequencing Data. PLoS Computational Biology, 12(6), 1–12. 10.1371/journal.pcbi.1004957

24. Jones, M. W., Abatzoglou, J. T., Veraverbeke, S., Andela, N., Lasslop, G., Forkel, M., Smith, A. J. P., Burton, C., Betts, R. A., van der Werf, G. R., Sitch, S., Canadell, J. G., Santín, C., Kolden, C., Doerr, S. H., & Le Quéré, C. (2022). Global and Regional Trends and Drivers of Fire Under Climate Change. Reviews of Geophysics, 60(3), 1–76. 10.1029/2020RG000726

25. Kelly, L. T., Giljohann, K. M., Duane, A., Aquilué, N., Archibald, S., Batllori, E., Bennett, A. F., Buckland, S. T., Canelles, Q., Clarke, M. F., Fortin, M. J., Hermoso, V., Herrando, S., Keane, R. E., Lake, F. K., McCarthy, M. A., Morán-Ordóñez, A., Parr, C. L., Pausas, J. G., … Brotons, L. (2020). Fire and biodiversity in the Anthropocene. Science, 370(6519). 10.1126/science.abb0355

26. Knicker, H. (2007). How does fire affect the nature and stability of soil organic nitrogen and carbon? A review. Biogeochemistry, 85(1), 91–118. 10.1007/s10533-007-9104-4

27. Kostenko, O., van de Voorde, T. F. J., Mulder, P. P. J., van der Putten, W. H., & Martijn Bezemer, T. (2012). Legacy effects of aboveground-belowground interactions. Ecology Letters, 15(8), 813–821. 10.1111/j.1461-0248.2012.01801.x

28. Langfelder, P., & Horvath, S. (2008). WGCNA: An R package for weighted correlation network analysis. BMC Bioinformatics, 9. 10.1186/1471-2105-9-559

29. Lasslop, G., Coppola, A. I., Voulgarakis, A., Yue, C., & Veraverbeke, S. (2019). Influence of Fire on the Carbon Cycle and Climate. Current Climate Change Reports, 5(2), 112–123. 10.1007/s40641-019-00128-9

30. Lemoine, G. G., Scott-Boyer, M. P., Ambroise, B., Périn, O., & Droit, A. (2021). GWENA: gene co-expression networks analysis and extended modules characterization in a single Bioconductor package. BMC Bioinformatics, 22(1), 1–20. 10.1186/s12859-021-04179-4

31. Li, J., Pei, J., Liu, J., Wu, J., Li, B., Fang, C., & Nie, M. (2021). Spatiotemporal variability of fire effects on soil carbon and nitrogen: A global meta-analysis. Global Change Biology, 27(17), 4196–4206. 10.1111/gcb.15742

32. Ling, L., Fu, Y., Jeewani, P. H., Tang, C., Pan, S., Reid, B. J., Gunina, A., Li, Y., Li, Y., Cai, Y., Kuzyakov, Y., Li, Y.Su, W. qin, Singh, B. P., Luo, Y., & Xu, J. (2021). Organic matter chemistry and bacterial community structure regulate decomposition processes in post-fire forest soils. Soil Biology and Biochemistry, 160, 108311. 10.1016/j.soilbio.2021.108311

33. Lopez, A. M., Avila, C. C. E., VanderRoest, J. P., Roth, H. K., Fendorf, S., & Borch, T. (2024). Molecular insights and impacts of wildfire-induced soil chemical changes. Nature Reviews Earth and Environment, 5(6), 431–446. 10.1038/s43017-024-00548-8

34. Lucas-Borja, M. E., Miralles, I., Ortega, R., Plaza-Álvarez, P. A., Gonzalez-Romero, J., Sagra, J., Soriano-Rodríguez, M., Certini, G., Moya, D., & Heras, J. (2019). Immediate fire-induced changes in soil microbial community composition in an outdoor experimental controlled system. Science of the Total Environment, 696. 10.1016/j.scitotenv.2019.134033

35. Ludwig, S. M., Alexander, H. D., Kielland, K., Mann, P. J., Natali, S. M., & Ruess, R. W. (2018). Fire severity effects on soil carbon and nutrients and microbial processes in a Siberian larch forest. Global Change Biology, 24(12), 5841–5852. 10.1111/gcb.14455

36. Martin, G., Morrissey, E. M., Carson, W., & Freedman, Z. B. (2022). A legacy of fire emerges from multiple disturbances to most shape microbial and nitrogen dynamics in a deciduous forest. Soil Biology and Biochemistry, 169, 108672.

37. McClure, R. S., Lee, J. Y., Chowdhury, T. R., Bottos, E. M., White, R. A., Kim, Y. M., Nicora, C. D., Metz, T. O., Hofmockel, K. S., Jansson, J. K., & Song, H. S. (2020). Integrated network modeling approach defines key metabolic responses of soil microbiomes to perturbations. Scientific Reports, 10(1). 10.1038/s41598-020-67878-7

38. Menges, E. S., Craddock, A., Salo, J., Zinthefer, R., & Weekley, C. W. (2008). Gap ecology in Florida scrub: Species occurrence, diversity and gap properties. Journal of Vegetation Science, 19(4), 503–514. 10.3170/2008-8-18399

39. Menges, E. S., Crate, S. J. H., & Quintana-Ascencio, P. F. (2017). Dynamics of gaps, vegetation, and plant species with and without fire. American Journal of Botany, 104(12), 1825–1836. 10.3732/ajb.1700175

40. Menges, E. S., Quintana-Ascencio, P. F., Sclater, V. L., Koontz, S. M., Smith, S. A., & David, A. S. (2018). Predicting landscape-level distribution and abundance: Integrating demography, fire, elevation and landscape habitat configuration. Journal of Ecology, 106(6), 2395–2408. 10.1111/1365-2745.12985

41. Miller, A. D., Inamine, H., Buckling, A., Roxburgh, S. H., & Shea, K. (2021). How disturbance history alters invasion success: biotic legacies and regime change. Ecology Letters, 24(4), 687–697. 10.1111/ele.13685

42. Miller, J. E. D., & Safford, H. D. (2020). Are plant community responses to wildfire contingent upon historical disturbance regimes? Global Ecology and Biogeography, 29(10), 1621– 1633. 10.1111/geb.13115

43. Morriën, E., Hannula, S. E., Snoek, L. B., Helmsing, N. R., Zweers, H., Hollander, M. De, Soto, R. L., Bouffaud, M. L., Buée, M., Dimmers, W., Duyts, H., Geisen, S., Girlanda, M., Jørgensen, H.-B., Jensen, J., Plassart, P., Redecker, D., Schmelz, R. M., Schmidt, O., … Putten, W. H. van der. (2016). Soil networks become more connected and take up more carbon as nature restoration progresses. Nature Communications, *in press*. 10.1038/ncomms14349

44. Müller, L. M., & Bahn, M. (2022). Drought legacies and ecosystem responses to subsequent drought. In Global Change Biology (Vol. 28, Issue 17, pp. 5086–5103). John Wiley and Sons Inc. 10.1111/gcb.16270

45. Muñoz-Rojas, M., Erickson, T. E., Martini, D., Dixon, K. W., & Merritt, D. J. (2016). Soil physicochemical and microbiological indicators of short, medium and long term post-fire recovery in semi-arid ecosystems. Ecological Indicators, 63, 14–22. 10.1016/j.ecolind.2015.11.038

46. Naylor, D., Sadler, N., Bhattacharjee, A., Graham, E. B., Anderton, C. R., McClure, R., Lipton, M., Hofmockel, K. S., & Jansson, J. K. (2020). Soil microbiomes under climate change and implications for carbon cycling. Annual Review of Environment and Resources, 45, 29–59. 10.1146/annurev-environ-012320-082720

47. Nelson, A. R., Narrowe, A. B., Rhoades, C. C., Fegel, T. S., Daly, R. A., Roth, H. K., Chu, R. K., Amundson, K. K., Young, R. B., Steindorff, A. S., Mondo, S. J., Grigoriev, I. V., Salamov, A., Borch, T., & Wilkins, M. J. (2022). Wildfire-dependent changes in soil microbiome diversity and function. Nature Microbiology, 7(9), 1419–1430. 10.1038/s41564-022-01203-y

48. Nelson, M. B., Martiny, A. C., & Martiny, J. B. H. (2016). Global biogeography of microbial nitrogen-cycling traits in soil. Proceedings of the National Academy of Sciences, 113(29), 8033–8040. 10.1073/pnas.1601070113

49. Pellegrini, A. F. A., Harden, J., Georgiou, K., Hemes, K. S., Malhotra, A., Nolan, C. J., & Jackson, R. B. (2022). Fire effects on the persistence of soil organic matter and long-term carbon storage. Nature Geoscience, 15(1), 5–13. 10.1038/s41561-021-00867-1

50. Pellegrini, A. F. A., Hobbie, S. E., Reich, P. B., Jumpponen, A., Brookshire, E. N. J., Caprio, A. C., Coetsee, C., & Jackson, R. B. (2020). Repeated fire shifts carbon and nitrogen cycling by changing plant inputs and soil decomposition across ecosystems. Ecological Monographs, 90(4),&lt;otherinfo> 1&lt;/otherinfo>–20. 10.1002/ecm.1409

51. Pellegrini, A. F. A., & Jackson, R. B. (2020). The long and short of it: A review of the timescales of how fire affects soils using the pulse-press framework. In Advances in Ecological Research (1st ed., Vol. 62). Elsevier Ltd. 10.1016/bs.aecr.2020.01.010

52. Pérez-Valera, E., Goberna, M., & Verdú, M. (2019). Fire modulates ecosystem functioning through the phylogenetic structure of soil bacterial communities. Soil Biology and Biochemistry, 129(October 2018), 80–89. 10.1016/j.soilbio.2018.11.007

53. Perring, M. P., De Frenne, P., Baeten, L., Maes, S. L., Depauw, L., Blondeel, H., Carón, M. M., & Verheyen, K. (2016). Global environmental change effects on ecosystems: The importance of land-use legacies. Global Change Biology, 22(4), 1361–1371. 10.1111/gcb.13146

54. Pierson, D. N., Robichaud, P. R., Rhoades, C. C., & Brown, R. E. (2019). Soil carbon and nitrogen eroded after severe wildfire and erosion mitigation treatments. International Journal of Wildland Fire, 28(10), 814–821. 10.1071/WF18193

55. Pingree, M. R. A., & DeLuca, T. H. (2017). Function of wildfire-deposited pyrogenic carbon in terrestrial ecosystems. In Frontiers in Environmental Science (Vol. 5, Issue AUG). Frontiers Media S.A. 10.3389/fenvs.2017.00053

56. Pressler, Y., Moore, J. C., & Cotrufo, M. F. (2019). Belowground community responses to fire: meta-analysis reveals contrasting responses of soil microorganisms and mesofauna. Oikos, 128(3), 309–327. 10.1111/oik.05738

57. Qu, X., Li, X., Bardgett, R. D., Kuzyakov, Y., Revillini, D., Sonne, C., Xia, C., Ruan, H., Liu, Y., Reich, P. B., & Delgado-Baquerizo, M. (2024). Deforestation impacts soil biodiversity and ecosystem services worldwide. Proceedings of the National Academy of Sciences, 121(13), e2318475121. 10.1073/pnas

58. R Core Team. (2020). R: A language and environment for statistical computing. (3.3.0). http://www.r-project.org/

59. Revillini, D., David, A. S., Menges, E. S., Main, K. N., Afkhami, M. E., & Searcy, C. A. (2021). Microbiome-mediated response to pulse fire disturbance outweighs the effects of fire legacy on plant performance. New Phytologist. 10.1111/nph.17689

60. Santín, C., & Doerr, S. H. (2016). Fire effects on soils: The human dimension. Philosophical Transactions of the Royal Society B: Biological Sciences, 371(1696), 28–34. 10.1098/rstb.2015.0171

61. Scharer, M., Andreas, L., & Kahmen, A. (2023). Post-drought compensatory growth in perennial grasslands is determined by legacy effects of the soil and not by plants. New Phytologist. 10.1111/nph.19291

62. Schimel, J. (2013). Soil carbon: Microbes and global carbon. Nature Publishing Group, 3(10), 867–868. 10.1038/nclimate2015

63. Singh, N., Abiven, S., Torn, M. S., & Schmidt, M. W. I. (2012). Fire-derived organic carbon in soil turns over on a centennial scale. Biogeosciences, 9(8), 2847–2857. 10.5194/bg-9-2847-2012

64. Stephens, S. L., Westerling, A. L. R., Hurteau, M. D., Peery, M. Z., Schultz, C. A., & Thompson, S. (2020). Fire and climate change: conserving seasonally dry forests is still possible. Frontiers in Ecology and the Environment, 18(6), 354–360. 10.1002/fee.2218

65. Subedi, S. C., Allen, P., Vidales, R., Sternberg, L., Ross, M., & Afkhami, M. E. (2022). Salinity legacy: Foliar microbiome’s history affects mutualist-conferred salinity tolerance. Ecology, 103(6), 1–14. 10.1002/ecy.3679

66. Tecon, R., & Or, D. (2017). Biophysical processes supporting the diversity of microbial life in soil. FEMS Microbiology Reviews, 41(5), 599–623. 10.1093/femsre/fux039

67. Wan, S., Hui, D., & Luo, Y. (2001). Fire Effects on Nitrogen Pools and Dynamics in Terrestrial Ecosystems: A Meta-Analysis. In Source: Ecological Applications (Vol. 11, Issue 5).

68. Wang, Q., Zhong, M., & Wang, S. (2012). A meta-analysis on the response of microbial biomass, dissolved organic matter, respiration, and N mineralization in mineral soil to fire in forest ecosystems. Forest Ecology and Management, 271, 91–97. 10.1016/j.foreco.2012.02.006

69. Westreich, S. T., Treiber, M. L., Mills, D. A., Korf, I., & Lemay, D. G. (2018). SAMSA2: A standalone metatranscriptome analysis pipeline. BMC Bioinformatics, 19(1). 10.1186/s12859-018-2189-z

70. Wubs, E. R. J., van der Putten, W. H., Mortimer, S. R., Korthals, G. W., Duyts, H., Wagenaar, R., & Bezemer, T. M. (2019). Single introductions of soil biota and plants generate long-term legacies in soil and plant community assembly. Ecology Letters, 22(7), 1145–1151. 10.1111/ele.13271

71. Yang, S., Zheng, Q., Yang, Y., Yuan, M., Ma, X., Chiariello, N. R., Docherty, K. M., Field, C. B., Gutknecht, J. L. M., Hungate, B. A., Niboyet, A., Le Roux, X., & Zhou, J. (2020). Fire affects the taxonomic and functional composition of soil microbial communities, with cascading effects on grassland ecosystem functioning. Global Change Biology, 26(2), 431–442. 10.1111/gcb.14852

72. Yu, G., Wang, L. G., Han, Y., & He, Q. Y. (2012). ClusterProfiler: An R package for comparing biological themes among gene clusters. OMICS A Journal of Integrative Biology, 16(5), 284–287. 10.1089/omi.2011.0118

