## Supplemental Information for "50-year fire legacy regulates soil microbial carbon and nutrient cycling responses to new fire"

**Supplemental Table 1**. Gap selection identifiers, fire treatment, fire legacy metrics for each sample, and sampling coordinates.

| **sample ID-seq** | **Fire treatment** | **Gap** | **TSF** | **Num_fires** | **Latitude** | **Longtitude** |
| --- | --- | --- | --- | --- | --- | --- |
| fire_15.merged.ribodepleted.fastq.nr | Burned | 40 | 1 | 2 | 27.18921022 | -81.36181089 |
| fire_13.merged.ribodepleted.fastq.nr | Burned | 13 | 2 | 3 | 27.17537323 | -81.36401738 |
| fire_19E.merged.ribodepleted.fastq.nr | Burned | 43 | 5 | 4 | 27.1784656 | -81.36220022 |
| fire_59.merged.ribodepleted.fastq.nr | Burned | 15 | 5 | 5 | 27.17832375 | -81.36119027 |
| fire_70N.merged.ribodepleted.fastq.nr | Burned | 19E | 7 | 5 | 27.17933675 | -81.36047678 |
| fire_31.merged.ribodepleted.fastq.nr | Burned | 59 | 9 | 3 | 27.16604849 | -81.36345216 |
| fire_94.merged.ribodepleted.fastq.nr | Burned | 94 | 9 | 2 | 27.1607344 | -81.36207234 |
| fire_49W.merged.ribodepleted.fastq.nr | Burned | 9 | 10 | 2 | 27.15608518 | -81.36193624 |
| fire_40.merged.ribodepleted.fastq.nr | Burned | 70N | 11 | 2 | 27.15253752 | -81.36524528 |
| fire_90.merged.ribodepleted.fastq.nr | Burned | 90 | 11 | 3 | 27.14842875 | -81.35859121 |
| fire_9.merged.ribodepleted.fastq.nr | Burned | 31 | 15 | 3 | 27.18256767 | -81.36443444 |
| fire_97.merged.ribodepleted.fastq.nr | Burned | 97 | 16 | 2 | 27.14492659 | -81.35826718 |
| fire_24N.merged.ribodepleted.fastq.nr | Burned | 49W | 18 | 2 | 27.1359164 | -81.3652518 |
| fire_8.merged.ribodepleted.fastq.nr | Burned | 24N | 18 | 4 | 27.19540535 | -81.36461752 |
| fire_43.merged.ribodepleted.fastq.nr | Burned | 8 | 21 | 3 | 27.13031115 | -81.36473914 |
| fire_93.merged.ribodepleted.fastq.nr | Burned | 93 | 33 | 2 | 27.12455281 | -81.36057342 |
| control_23.merged.ribodepleted.fastq.nr | Unburned | 35N | 1 | 3 | 27.19151178 | -81.35854717 |
| control_24N.merged.ribodepleted.fastq.nr | Unburned | 40 | 1 | 2 | 27.12394072 | -81.35985506 |
| control_13.merged.ribodepleted.fastq.nr | Unburned | 13 | 2 | 3 | 27.13715379 | -81.36505399 |
| control_43.merged.ribodepleted.fastq.nr | Unburned | 5E | 3 | 3 | 27.20744112 | -81.35716427 |
| control_28.merged.ribodepleted.fastq.nr | Unburned | 43 | 5 | 4 | 27.18921022 | -81.36181089 |
| control_46W.merged.ribodepleted.fastq.nr | Unburned | 20 | 7 | 4 | 27.18648957 | -81.35598699 |
| control_89.merged.ribodepleted.fastq.nr | Unburned | 22E | 7 | 7 | 27.17802492 | -81.36502538 |
| control_40.merged.ribodepleted.fastq.nr | Unburned | 59 | 9 | 3 | 27.17933675 | -81.36047678 |
| control_45S.merged.ribodepleted.fastq.nr | Unburned | 65E | 9 | 4 | 27.1607344 | -81.36207234 |
| control_94.merged.ribodepleted.fastq.nr | Unburned | 94 | 9 | 2 | 27.15253752 | -81.36524528 |
| control_20.merged.ribodepleted.fastq.nr | Unburned | 28 | 10 | 5 | 27.14842875 | -81.35859121 |
| control_65E.merged.ribodepleted.fastq.nr | Unburned | 9 | 10 | 2 | 27.14074018 | -81.36422804 |
| control_59.merged.ribodepleted.fastq.nr | Unburned | 89 | 11 | 4 | 27.1359164 | -81.3652518 |
| control_90.merged.ribodepleted.fastq.nr | Unburned | 90 | 11 | 3 | 27.13061204 | -81.35655658 |
| control_22E.merged.ribodepleted.fastq.nr | Unburned | 31 | 15 | 3 | 27.19256805 | -81.35856064 |
| control_99.merged.ribodepleted.fastq.nr | Unburned | 99 | 16 | 1 | 27.19151178 | -81.35854717 |
| control_31.merged.ribodepleted.fastq.nr | Unburned | 45S | 18 | 2 | 27.12394072 | -81.35985506 |
| control_5E.merged.ribodepleted.fastq.nr | Unburned | 23 | 18 | 5 | 27.12137141 | -81.36190517 |
| control_9.merged.ribodepleted.fastq.nr | Unburned | 24N | 18 | 4 | 27.13715379 | -81.36505399 |
| control_37.merged.ribodepleted.fastq.nr | Unburned | 46W | 24 | 2 | 27.20769755 | -81.35573123 |

**Supplemental Table 2.** Carbon, nitrogen, and phosphorus cycling gene list used to identify microbial KEGG genes associated with critical carbon and nutrient cycling functions.

| **Gene** | **Enzyme** | **KEGG ortholog** | **Function** | **Function specific** | **EC** |
| --- | --- | --- | --- | --- | --- |
| *ACADM, acd* | acyl-CoA dehydrogenase | K00249 | Fatty acid degradation |  | EC:1.3.8.7 |
| *ACADVL* | very long chain acyl-CoA dehydrogenase | K09479 | Fatty acid degradation |  | EC:1.3.8.9 |
| *accD6* | acetyl-CoA/propionyl-CoA carboxylase carboxyl transferase subunit | K18472 | Carbohydrate metabolism | Pyruvate metabolism | EC:6.4.1.2 ; 6.4.1.3 ; 2.1.3.15 |
| *acdA* | acetate---CoA ligase (ADP-forming) subunit alpha | K01905 | Carbohydrate metabolism | Glucolysis | EC:6.2.1.13 |
| *acdB* | acetate---CoA ligase (ADP-forming) subunit beta | K22224 | Carbohydrate metabolism | Glucolysis | EC:6.2.1.13 |
| *ACMSD* | aminocarboxymuconate-semialdehyde decarboxylase | K03392 | Aromatic aminoacid metabolism | Tryptophan metabolism | EC:4.1.1.45 |
| *ACSL, fadD* | long-chain acyl-CoA synthetase | K01897 | Fatty acid biosynthesis |  | EC:6.2.1.3 |
| *adh* | alcohol dehydrogenase | K00001 | Carbohydrate metabolism | Glycolysis | EC:1.1.1.1 |
| *aldB* | aldehyde dehydrogenase | K00138 | Carbohydrate metabolism | Glucolysis | EC:1.2.1.- |
| *ALDH* | aldehyde dehydrogenase (NAD+) | K00128 | Carbohydrate metabolism | Glucolysis | EC:1.2.1.3 |
| *ALDH3* | aldehyde dehydrogenase (NAD(P)+) | K00129 | Carbohydrate metabolism | Glucolysis | EC:1.2.1.5 |
| *amiE* | amidase | K01426 | Aromatic aminoacid metabolism | Aminobenzoate degradation | EC:3.5.1.4 |
| *AOC3, AOC2, tynA* | primary-amine oxidase | K00276 | Aromatic aminoacid metabolism | Glycine, serine and threonine metabolism | EC:1.4.3.21 |
| *aspB* | aspartate aminotransferase | K00812 | Aromatic aminoacid metabolism | Alanine, aspartate and glutamate metabolism | EC:2.6.1.1 |
| *benA-xylX* | benzoate/toluate 1,2-dioxygenase subunit alpha | K05549 | Aromatic compounds degradation | Benzoate degradation | EC:1.14.12.10 ; 1.14.12.- |
| *benD-xylL* | dihydroxycyclohexadiene carboxylate dehydrogenase | K05783 | Aromatic compounds degradation | Benzoate degradation | EC:1.3.1.25 ; 1.3.1.- |
| *bgaB, lacA* | beta-galactosidase | K12308 | Carbohydrate metabolism | Hemicellulose-degrading | EC:3.2.1.23 |
| *bglB* | beta-glucosidase | K05350 | Carbohydrate metabolism | Hemicellulose-degrading | EC:3.2.1.21 |
| *bglX* | beta-glucosidase | K05349 | Carbohydrate metabolism | Hemicellulose-degrading | EC:3.2.1.21 |
| *catA* | catechol 1,2-dioxygenase | K03381 | Aromatic compounds degradation | Benzoate degradation | EC:1.13.11.1 |
| *catC* | muconolactone D-isomerase | K03464 | Aromatic compounds degradation | Benzoate degradation | EC:5.3.3.4 |
| *catE* | catechol 2,3-dioxygenase | K07104 | Aromatic compounds degradation | Benzoate degradation | EC:1.13.11.2 |
| *crt* | enoyl-CoA hydratase | K01715 | Carbohydrate metabolism | Butanoate metabolism | EC:4.2.1.17 |
| *CTH* | cystathionine gamma-lyase | K01758 | Sulfur aminoacid metabolism | Cysteine and methionine metabolism | EC:4.4.1.1 |
| *CYP71D18* | (S)-limonene 6-monooxygenase | K07381 | Aromatic compounds degradation | Benzoate degradation | EC:1.14.14.51 |
| *cypD_E, CYP102A, CYP505* | cytochrome P450 / NADPH-cytochrome P450 reductase | K14338 | Fatty acid degradation | Aminobenzoate degradation | EC:1.14.14.1 ; 1.6.2.4 |
| *DBT, bkdB* | 2-oxoisovalerate dehydrogenase E2 component (dihydrolipoyl transacylase) | K09699 | Amino acid metabolism | Leucine degradation | EC:2.3.1.168 |
| *DDC, TDC* | aromatic-L-amino-acid/L-tryptophan decarboxylase | K01593 | Aromatic aminoacid metabolism | Tyrosine metabolism | EC:4.1.1.28 ; 4.1.1.105 |
| *deoB* | phosphopentomutase | K01839 | Phosphorus metabolism | Pentose phosphate pathway | EC:5.4.2.7 |
| *desA2* | acyl-[acyl-carrier-protein] desaturase | K03922 | Fatty acid biosynthesis |  | EC:1.14.19.2 |
| *dmpB, xylE* | catechol 2,3-dioxygenase | K00446 | Aromatic compounds degradation | Benzoate degradation | EC:1.13.11.2 |
| *dmpC, xylG, praB* | aminomuconate-semialdehyde/2-hydroxymuconate-6-semialdehyde dehydrogenase | K10217 | Aromatic compounds degradation | Benzoate degradation | EC:1.2.1.32 ; 1.2.1.85 |
| *edd* | phosphogluconate dehydratase | K01690 | Phosphorus metabolism | Pentose phosphate pathway | EC:4.2.1.12 |
| *ENO, eno* | enolase | K01689 | Carbohydrate metabolism | Glucolysis | EC:4.2.1.11 |
| *FAB2, SSI2,* | desA1 acyl-[acyl-carrier-protein] desaturase | K03921 | Fatty acid biosynthesis |  | EC:1.14.19.2 ; 1.14.19.11 ; 1.14.19.26 |
| *fabH* | 3-oxoacyl-[acyl-carrier-protein] synthase III | K00648 | Fatty acid biosynthesis |  | EC:2.3.1.180 |
| *fabV, ter* | enoyl-[acyl-carrier protein] reductase / trans-2-enoyl-CoA reductase (NAD+) | K00209 | Carbohydrate metabolism | Butanoate metabolism | EC:1.3.1.9 ; 1.3.1.44 |
| *fadA, fadI* | acetyl-CoA acyltransferase | K00632 | Fatty acid degradation | Benzoate degradation | EC:2.3.1.16 |
| *fadE* | acyl-CoA dehydrogenase | K06445 | Fatty acid degradation |  | EC:1.3.99.- |
| *fadJ* | 3-hydroxyacyl-CoA dehydrogenase / enoyl-CoA hydratase/ 3-hydroxybutyryl-CoA epimerase | K01782 | Fatty acid degradation |  | EC:1.1.1.35 ; 4.2.1.17 ; 5.1.2.3 |
| *fas* | fatty acid synthase, bacteria type | K11533 | Fatty acid biosynthesis |  | EC:2.3.1.- |
| *FAS1* | fatty acid synthase subunit beta, fungi type | K00668 | Fatty acid biosynthesis |  | EC:2.3.1.86 |
| *FASN* | fatty acid synthase, animal type | K00665 | Fatty acid biosynthesis |  | EC:2.3.1.85 |
| *fbaB* | fructose-bisphosphate aldolase, class I | K11645 | Carbohydrate metabolism | Glycolysis | EC:4.1.2.13 |
| *fumA* | fumarate hydratase subunit alpha | K01677 | Carbohydrate metabolism | Citrate cycling | EC:4.2.1.2 |
| *GBE1, glgB* | 1,4-alpha-glucan branching enzyme | K00700 | Carbohydrate metabolism | Starch and sucrose | EC:2.4.1.18 |
| *gdh* | glucose 1-dehydrogenase | K00034 | Phosphorus metabolism | Pentose phosphate pathway | EC:1.1.1.47 |
| *ggt* | gamma-glutamyltranspeptidase / glutathione hydrolase | K00681 | Sulfur aminoacid metabolism | Taurine and hypotaurine metabolism | EC:2.3.2.2 ; 3.4.19.13 |
| *glgC* | glucose-1-phosphate adenylyltransferase | K00975 | Carbohydrate metabolism | Starch and sucrose | EC:2.7.7.27 |
| *glgM* | alpha-maltose-1-phosphate synthase | K16148 | Carbohydrate metabolism | Starch and sucrose | EC:2.4.1.342 |
| *glk* | glucokinase | K00845 | Carbohydrate metabolism | Starch and sucrose | EC:2.7.1.2 |
| *glpQ, ugpQ* | glycerophosphoryl diester phosphodiesterase | K01126 | Phosphorus metabolism | Inorganic P solubilization | EC:3.1.4.46 |
| *gnl, RGN* | gluconolactonase | K01053 | Phosphorus metabolism | Pentose phosphate pathway | EC:3.1.1.17 |
| *HGD, hmgA* | homogentisate 1,2-dioxygenase | K00451 | Aromatic aminoacid metabolism | Tyrosine metabolism | EC:1.13.11.5 |
| *hisC* | histidinol-phosphate aminotransferase | K00817 | Aromatic aminoacid metabolism | Histidine metabolism | EC:2.6.1.9 |
| *hpaB* | 4-hydroxyphenylacetate 3-monooxygenase | K00483 | Aromatic aminoacid metabolism | Tyrosine metabolism | EC:1.14.14.9 |
| *HPD, hppD* | 4-hydroxyphenylpyruvate dioxygenase | K00457 | Aromatic aminoacid metabolism | Tyrosine metabolism | EC:1.13.11.27 |
| *hxlB* | 6-phospho-3-hexuloisomerase | K08094 | Phosphorus metabolism | Pentose phosphate pathway | EC:5.3.1.27 |
| *IDH1, IDH2, icd* | isocitrate dehydrogenase | K00031 | Carbohydrate metabolism | Citrate cycling | EC:1.1.1.42 |
| *ilvA, tdcB* | threonine dehydratase | K01754 | Amino acid metabolism | Leucine degradation | EC:4.3.1.19 |
| *ilvB, ilvG, ilvI* | acetolactate synthase I/II/III large subunit | K01652 | Amino acid metabolism | Valine biosynthesis | EC:2.2.1.6 |
| *ilvC* | ketol-acid reductoisomerase | K00053 | Amino acid metabolism | Valine biosynthesis | EC:1.1.1.86 |
| *ilvD* | dihydroxy-acid dehydratase | K01687 | Amino acid metabolism | Valine biosynthesis | EC:4.2.1.9 |
| *ilvE* | branched-chain amino acid aminotransferase | K00826 | Sulfur aminoacid metabolism | Cysteine and methionine metabolism | EC:2.6.1.42 |
| *ilvH, ilvN* | acetolactate synthase I/III small subunit | K01653 | Amino acid metabolism | Valine biosynthesis | EC:2.2.1.6 |
| *IMA, malL* | oligo-1,6-glucosidase | K01182 | Carbohydrate metabolism | Starch and sucrose | EC:3.2.1.10 |
| *INV, sacA* | beta-fructofuranosidase | K01193 | Carbohydrate metabolism | Starch and sucrose | EC:3.2.1.26 |
| *IVD, ivd* | isovaleryl-CoA dehydrogenase | K00253 | Amino acid metabolism | Leucine degradation | EC:1.3.8.4 |
| *K01622* | fructose 1,6-bisphosphate aldolase/phosphatase | K01622 | Phosphorus metabolism | Pentose phosphate pathway | EC:4.1.2.13 ; 3.1.3.11 |
| *K03822* | putative long chain acyl-CoA synthase | K03822 | Lipid biosynthesis | Lipid biosynthesis | EC:6.2.1.- |
| *K16164* | acylpyruvate hydrolase | K16164 | Aromatic aminoacid metabolism | Tyrosine metabolism | EC:3.7.1.5 |
| *katG* | catalase-peroxidase | K03782 | Aromatic aminoacid metabolism | Phenylalanine metabolism | EC:1.11.1.21 |
| *korB, oorB, oforB* | 2-oxoglutarate/2-oxoacid ferredoxin oxidoreductase subunit beta | K00175 | Carbohydrate metabolism | Glucolysis | EC:1.2.7.3 ; 1.2.7.11 |
| *kynB* | arylformamidase | K07130 | Aromatic aminoacid metabolism | Tryptophan metabolism | EC:3.5.1.9 |
| *liuC* | methylglutaconyl-CoA hydratase | K13766 | Amino acid metabolism | Leucine degradation | EC:4.2.1.18 |
| *lpxC-fabZ* | UDP-3-O-[3-hydroxymyristoyl] N-acetylglucosamine deacetylase / 3-hydroxyacyl-[acyl-carrier-protein] dehydratase | K16363 | Fatty acid biosynthesis | Lipopolysaccharide biosynthesis | EC:3.5.1.108 ; 4.2.1.59 |
| *maeB* | malate dehydrogenase (oxaloacetate-decarboxylating)(NADP+) | K00029 | Carbohydrate metabolism | Pyruvate metabolism | EC:1.1.1.40 |
| *malZ* | alpha-glucosidase | K01187 | Carbohydrate metabolism | Hemicellulose-degrading | EC:3.2.1.20 |
| *MAO, aofH* | monoamine oxidase | K00274 | Aromatic aminoacid metabolism | Glycine, serine and threonine metabolism | EC:1.4.3.4 |
| *mcmA1* | methylmalonyl-CoA mutase, N-terminal domain | K01848 | Carbohydrate metabolism | Glyoxylate and dicarboxylate pathway | EC:5.4.99.2 |
| *metE* | 5-methyltetrahydropteroyltriglutamate--homocysteine methyltransferase | K00549 | Sulfur aminoacid metabolism | Cysteine and methionine metabolism | EC:2.1.1.14 |
| *metH, MTR* | 5-methyltetrahydrofolate--homocysteine methyltransferase | K00548 | Sulfur aminoacid metabolism | Cysteine and methionine metabolism | EC:2.1.1.13 |
| *metY* | O-acetylhomoserine (thiol)-lyase | K01740 | Sulfur aminoacid metabolism | Cysteine and methionine metabolism | EC:2.5.1.49 |
| *mqo* | malate dehydrogenase (quinone) | K00116 | Carbohydrate metabolism | Citrate cycling | EC:1.1.5.4 |
| *mtaD* | 5-methylthioadenosine/S-adenosylhomocysteine deaminase | K12960 | Sulfur aminoacid metabolism | Cysteine and methionine metabolism | EC:3.5.4.31 ; 3.5.4.28 |
| *nagK* | fumarylpyruvate hydrolase | K16165 | Aromatic aminoacid metabolism | Tyrosine metabolism | EC:3.7.1.20 |
| *narH, narY, nxrB* | nitrate reductase / nitrite oxidoreductase, beta subunit | K00371 | Nitrogen metabolism | Nitrate metabolism | EC:1.7.5.1 ; 1.7.99.- |
| *nifB* | nitrogen fixation protein | K02585 | Nitrogen metabolism | Nitrogen-fixation | EC:NA |
| *nifH* | nitrogenase iron protein | K02588 | Nitrogen metabolism | Nitrogen-fixation | EC:NA |
| *nifK* | nitrogenase molybdenum-iron protein beta chain | K02591 | Nitrogen metabolism | Nitrogen-fixation | EC:1.18.6.1 |
| *nirA* | ferredoxin-nitrite reductase | K00366 | Nitrogen metabolism | Nitrite metabolism | EC:1.7.7.1 |
| *nirB* | nitrite reductase (NADH) large subunit | K00362 | Nitrogen metabolism | Nitrite metabolism | EC:1.7.1.15 |
| *nirD* | nitrite reductase (NADH) small subunit | K00363 | Nitrogen metabolism | Nitrite metabolism | EC:1.7.1.15 |
| *nirK* | nitrite reductase (NO-forming) | K00368 | Nitrogen metabolism | Nitrite metabolism | EC:1.7.2.1 |
| *nirS* | nitrite reductase (NO-forming) / hydroxylamine reductase | K15864 | Nitrogen metabolism | Nitrite metabolism | EC:1.7.2.1 ; 1.7.99.1 |
| *norQ* | nitric oxide reductase NorQ protein | K04748 | Nitrogen metabolism | Nitric oxide-reduction | EC:NA |
| *NRT2, narK, nrtP, nasA* | MFS transporter, NNP family, nitrate/nitrite transporter | K02575 | Nitrogen metabolism | N-transport | EC:NA |
| *paaF, echA* | enoyl-CoA hydratase | K01692 | Fatty acid degradation |  | EC:4.2.1.17 |
| *paaH, hbd, fadB, mmgB* | 3-hydroxybutyryl-CoA dehydrogenase | K00074 | Aromatic aminoacid metabolism | Butanoate metabolism | EC:1.1.1.157 |
| *paaK* | phenylacetate-CoA ligase | K01912 | Aromatic aminoacid metabolism | Phenylalanine metabolism | EC:6.2.1.30 |
| *pcaB* | 3-carboxy-cis,cis-muconate cycloisomerase | K01857 | Aromatic compounds degradation | Protocatechuate dioxygenase | EC:5.5.1.2 |
| *pcaC* | 4-carboxymuconolactone decarboxylase | K01607 | Aromatic compounds degradation | Protocatechuate dioxygenase | EC:4.1.1.44 |
| *pcaD* | 3-oxoadipate enol-lactonase | K01055 | Aromatic compounds degradation | Protocatechuate dioxygenase | EC:3.1.1.24 |
| *pcaF* | 3-oxoadipyl-CoA thiolase | K07823 | Aromatic compounds degradation | Protocatechuate dioxygenase | EC:2.3.1.174 |
| *pcaG* | protocatechuate 3,4-dioxygenase, alpha subunit | K00448 | Aromatic compounds degradation | Protocatechuate dioxygenase | EC:1.13.11.3 |
| *pcaH* | protocatechuate 3,4-dioxygenase, beta subunit | K00449 | Aromatic compounds degradation | Protocatechuate dioxygenase | EC:1.13.11.3 |
| *pcaI* | 3-oxoadipate CoA-transferase, alpha subunit | K01031 | Aromatic compounds degradation | Protocatechuate dioxygenase | EC:2.8.3.6 |
| *pcaJ* | 3-oxoadipate CoA-transferase, beta subunit | K01032 | Aromatic compounds degradation | Protocatechuate dioxygenase | EC:2.8.3.6 |
| *pcaL* | 3-oxoadipate enol-lactonase/ 4-carboxymuconolactone decarboxylase | K14727 | Aromatic compounds degradation | Protocatechuate dioxygenase | EC:3.1.1.24 ; 4.1.1.44 |
| *pckA* | phosphoenolpyruvate carboxykinase (ATP) | K01610 | Carbohydrate metabolism | Glycolysis | EC:4.1.1.49 |
| *pckA, PCK* | phosphoenolpyruvate carboxykinase (GTP) | K01596 | Carbohydrate metabolism | Glucolysis | EC:4.1.1.32 |
| *pfkA, PFK* | 6-phosphofructokinase1 | K00850 | Carbohydrate metabolism | Glucolysis | EC:2.7.1.11 |
| *pgmB* | beta-phosphoglucomutase | K01838 | Carbohydrate metabolism | Starch and sucrose | EC:5.4.2.6 |
| *phnC* | phosphonate transport system ATP-binding protein | K02041 | Phosphorus metabolism | Organic P solubilization | EC:7.3.2.2 |
| *phnD* | phosphonate transport system substrate-binding protein | K02044 | Phosphorus metabolism | Organic P solubilization | EC:NA |
| *phnE* | phosphonate transport system permease protein | K02042 | Phosphorus metabolism | Organic P solubilization | EC:NA |
| *phnJ* | alpha-D-ribose 1-methylphosphonate 5-phosphate C-P lyase | K06163 | Phosphorus metabolism | Organic P solubilization | EC:4.7.1.1 |
| *PHO* | acid phosphatase | K01078 | Phosphorus metabolism | Inorganic P solubilization | EC:3.1.3.2 |
| *phoB* | two-component system, OmpR family, phosphate regulon response regulator | K07657 | Phosphorus metabolism | Phosphate transport | EC:NA |
| *phoD* | alkaline phosphatase D | K01113 | Phosphorus metabolism | Inorganic P solubilization | EC:3.1.3.1 |
| *phoR* | two-component system, OmpR family, phosphate regulon sensor histidine kinase | K07636 | Phosphorus metabolism | Phosphate transport | EC:2.7.13.3 |
| *phoU* | phosphate transport system protein | K02039 | Phosphorus metabolism | Phosphate transport | EC:NA |
| *pht3* | phthalate 4,5-dioxygenase | K18068 | Aromatic compounds degradation | Polycyclic aromatic hydrocarbon degradation | EC:1.14.12.7 |
| *pht4* | phthalate 4,5-cis-dihydrodiol dehydrogenase | K18067 | Aromatic compounds degradation | Polycyclic aromatic hydrocarbon degradation | EC:1.3.1.64 |
| *pmm-pgm* | phosphomannomutase / phosphoglucomutase | K15778 | Carbohydrate metabolism | Glucolysis | EC:5.4.2.8 ; 5.4.2.2 |
| *por, nifJ* | pyruvate-ferredoxin/flavodoxin oxidoreductase | K03737 | Carbohydrate metabolism | Pyruvate metabolism | EC:1.2.7.1 ; 1.2.7.- |
| *porA* | pyruvate ferredoxin oxidoreductase alpha subunit | K00169 | Carbohydrate metabolism | Pyruvate metabolism | EC:1.2.7.1 |
| *porB* | pyruvate ferredoxin oxidoreductase beta subunit | K00170 | Carbohydrate metabolism | Pyruvate metabolism | EC:1.2.7.1 |
| *porD* | pyruvate ferredoxin oxidoreductase delta subunit | K00171 | Carbohydrate metabolism | Pyruvate metabolism | EC:1.2.7.1 |
| *ppa* | inorganic pyrophosphatase | K01507 | Phosphorus metabolism | Inorganic P solubilization | EC:3.6.1.1 |
| *pps, ppsA* | pyruvate, water dikinase | K01007 | Carbohydrate metabolism | Pyruvate metabolism | EC:2.7.9.2 |
| *ppx-gppA* | exopolyphosphatase / guanosine-5'-triphosphate,3'-diphosphate pyrophosphatase | K01524 | Phosphorus metabolism | Inorganic P solubilization | EC:3.6.1.11 ; 3.6.1.40 |
| *pstA* | phosphate transport system permease protein | K02038 | Phosphorus metabolism | Phosphate transport | EC:NA |
| *pstB* | phosphate transport system ATP-binding protein | K02036 | Phosphorus metabolism | Phosphate transport | EC:7.3.2.1 |
| *pstC* | phosphate transport system permease protein | K02037 | Phosphorus metabolism | Phosphate transport | EC:NA |
| *pstS* | phosphate transport system substrate-binding protein | K02040 | Phosphorus metabolism | Phosphate transport | EC:NA |
| *PYG, glgP* | glycogen phosphorylase | K00688 | Carbohydrate metabolism | Starch and sucrose | EC:2.4.1.1 |
| *rbcL, cbbL* | ribulose-bisphosphate carboxylase large chain | K01601 | Carbohydrate metabolism | Glyoxylate and dicarboxylate pathway | EC:4.1.1.39 |
| *rbcS, cbbS* | ribulose-bisphosphate carboxylase small chain | K01602 | Carbohydrate metabolism | Glyoxylate and dicarboxylate pathway | EC:4.1.1.39 |
| *rbsK, RBKS* | ribokinase | K00852 | Phosphorus metabolism | Pentose phosphate pathway | EC:2.7.1.15 |
| *rpiB* | ribose 5-phosphate isomerase B | K01808 | Carbohydrate metabolism | Pentose phosphate pathway | EC:5.3.1.6 |
| *scrKscrK* | fructokinase | K00847 | Carbohydrate metabolism | Starch and sucrose | EC:2.7.1.4 |
| *sdhB, frdB* | succinate dehydrogenase iron-sulfur subunit | K00240 | Carbohydrate metabolism | Citrate cycling | EC:1.3.5.1 |
| *tal-pgi* | transaldolase / glucose-6-phosphate isomerase | K13810 | Carbohydrate metabolism | Glucolysis | EC:2.2.1.2 ; 5.3.1.9 |
| *tauD* | taurine dioxygenase | K03119 | Sulfur aminoacid metabolism | Cysteine and methionine metabolism | EC:1.14.11.17 |
| *TC.PIT* | inorganic phosphate transporter, PiT family | K03306 | Phosphorus metabolism | Phosphate transport | EC:NA |
| *tktA, tktB* | transketolase | K00615 | Carbohydrate metabolism | Pentose phosphate pathway | EC:2.2.1.1 |
| *TREH, treA, treF* | alpha,alpha-trehalase | K01194 | Carbohydrate metabolism | Starch and sucrose | EC:3.2.1.28 |
| *treY, glgY* | (1->4)-alpha-D-glucan 1-alpha-D-glucosylmutase | K06044 | Carbohydrate metabolism | Starch and sucrose | EC:5.4.99.15 |
| *UGP2, galU, galF* | UTP--glucose-1-phosphate uridylyltransferase | K00963 | Carbohydrate metabolism | Starch and sucrose | EC:2.7.7.9 |
| *ugpA* | sn-glycerol 3-phosphate transport system permease protein | K05814 | Phosphorus metabolism | Phosphate transport | EC:NA |
| *ugpC* | sn-glycerol 3-phosphate transport system ATP-binding protein | K05816 | Phosphorus metabolism | Phosphate transport | EC:7.6.2.10 |
| *URE* | urease | K01427 | Nitrogen metabolism | Urea-degradation | EC:3.5.1.5 |
| *ureA* | urease subunit gamma | K01430 | Nitrogen metabolism | Urea-degradation | EC:3.5.1.5 |
| *ureB* | urease subunit beta | K01429 | Nitrogen metabolism | Urea-degradation | EC:3.5.1.5 |
| *ureC* | urease subunit alpha | K01428 | Nitrogen metabolism | Urea-degradation | EC:3.5.1.5 |
| *vnfD* | vanadium-dependent nitrogenase alpha chain | K22896 | Nitrogen metabolism | Vanadium dependent nitrogen-fixation | EC:1.18.6.2 |
| *vnfG* | vanadium nitrogenase delta subunit | K22898 | Nitrogen metabolism | Vanadium dependent nitrogen-fixation | EC:1.18.6.2 |
| *vnfK* | vanadium-dependent nitrogenase beta chain | K22897 | Nitrogen metabolism | Vanadium dependent nitrogen-fixation | EC:1.18.6.2 |
| *xfp, xpk* | xylulose-5-phosphate/fructose-6-phosphate phosphoketolase | K01621 | Carbohydrate metabolism | Pentose phosphate pathway | EC:4.1.2.9 ; 4.1.2.22 |
| *xylC* | benzaldehyde dehydrogenase (NAD) | K00141 | Aromatic compounds degradation | Toluene degradation | EC:1.2.1.28 |
| *xynA* | endo-1,4-beta-xylanase | K01181 | Carbohydrate metabolism | Cellulose | EC:3.2.1.8 |

**Supplemental Table 3**. Significant results from linear mixed models for functional genes and metabolically active microbial taxa, including ANOVA type-III *P*-values.

| **Functions** |  |  |  |  |  |  |  |  |
| --- | --- | --- | --- | --- | --- | --- | --- | --- |
| **KEGG Ortholog** | **Gene** | **Model term** | **LMM estimate** | **SE** | **t-statistic** | **df** | **P value (LMM)** | **P value (ANOVA - Type III)** |
| K00053 | ilvC | Fire | -0.204 | 0.090 | -2.266 | 10.535 | 0.046 | 0.023 |
| K00138 | aldB | Fire ~ TSF | 0.031 | 0.010 | 3.231 | 10.906 | 0.008 | 0.001 |
| K00138 | aldB | TSF | -0.027 | 0.012 | -2.323 | 17.921 | 0.032 | 0.020 |
| K00141 | xylC | Fire ~ TSF ~ Num. fires | 0.008 | 0.003 | 2.658 | 8.963 | 0.026 | 0.008 |
| K00362 | nirB | Fire ~ TSF | -0.037 | 0.012 | -3.107 | 12.005 | 0.009 | 0.002 |
| K00362 | nirB | Fire ~ TSF ~ Num. fires | -0.034 | 0.012 | -2.911 | 18.867 | 0.009 | 0.004 |
| K00371 | narH | Fire | -0.321 | 0.072 | -4.433 | 13.241 | 0.001 | 0.000 |
| K00371 | narH | Num. fires | -0.317 | 0.077 | -4.122 | 25.585 | 0.000 | 0.000 |
| K00548 | metH | Fire ~ TSF ~ Num. fires | -0.050 | 0.023 | -2.147 | 22.970 | 0.043 | 0.032 |
| K00548 | metH | TSF | -0.062 | 0.028 | -2.220 | 21.564 | 0.037 | 0.026 |
| K00975 | glgC | Fire ~ TSF ~ Num. fires | -0.039 | 0.018 | -2.211 | 20.846 | 0.038 | 0.027 |
| K01007 | ppsA | Fire | -0.164 | 0.072 | -2.272 | 8.939 | 0.049 | 0.023 |
| K01187 | malZ | TSF | -0.048 | 0.022 | -2.145 | 19.321 | 0.045 | 0.032 |
| K01426 | amiE | Fire ~ TSF ~ Num. fires | -0.049 | 0.022 | -2.281 | 23.036 | 0.032 | 0.023 |
| K01426 | amiE | TSF | -0.051 | 0.026 | -1.998 | 21.235 | 0.059 | 0.046 |
| K01430 | ureA | Fire | 0.102 | 0.046 | 2.206 | 16.067 | 0.042 | 0.027 |
| K01524 | ppx-gppA | TSF ~ Num. fires | -0.038 | 0.017 | -2.204 | 28.000 | 0.036 | 0.028 |
| K01607 | pcaC | Fire ~ TSF | -0.037 | 0.018 | -2.071 | 13.501 | 0.058 | 0.038 |
| K01607 | pcaC | Fire ~ TSF ~ Num. fires | -0.041 | 0.018 | -2.311 | 20.004 | 0.032 | 0.021 |
| K01689 | eno | Fire | -0.366 | 0.125 | -2.930 | 28.000 | 0.007 | 0.003 |
| K01754 | ilvA/tdcB | Fire ~ TSF | 0.033 | 0.014 | 2.397 | 14.421 | 0.031 | 0.017 |
| K01808 | rpiB | Fire ~ Num. fires | -0.095 | 0.044 | -2.185 | 25.603 | 0.038 | 0.029 |
| K01808 | rpiB | Fire ~ TSF | 0.015 | 0.008 | 1.974 | 20.475 | 0.062 | 0.048 |
| K01897 | ACSL | TSF | -0.067 | 0.030 | -2.228 | 22.559 | 0.036 | 0.026 |
| K01905 | acdA | Fire ~ Num. fires | -0.103 | 0.037 | -2.819 | 9.593 | 0.019 | 0.005 |
| K01905 | acdA | Fire ~ TSF | 0.015 | 0.005 | 2.867 | 6.590 | 0.026 | 0.004 |
| K01905 | acdA | Fire ~ TSF ~ Num. fires | 0.019 | 0.006 | 3.321 | 9.535 | 0.008 | 0.001 |
| K01912 | paaK | Fire | -0.224 | 0.100 | -2.247 | 10.344 | 0.048 | 0.025 |
| K01912 | paaK | Fire ~ TSF ~ Num. fires | -0.035 | 0.016 | -2.206 | 18.865 | 0.040 | 0.027 |
| K02036 | pstB | Fire | -0.219 | 0.104 | -2.101 | 14.476 | 0.054 | 0.036 |
| K06445 | fadE | Num. fires | -0.114 | 0.054 | -2.121 | 24.407 | 0.044 | 0.034 |
| K06445 | fadE | TSF | -0.021 | 0.011 | -2.017 | 21.265 | 0.057 | 0.044 |
| K07130 | kynB | Fire | -0.157 | 0.060 | -2.624 | 15.971 | 0.018 | 0.009 |
| K07823 | pcaF | Fire | 0.082 | 0.032 | 2.540 | 20.128 | 0.019 | 0.011 |
| K07823 | pcaF | Num. fires | 0.062 | 0.030 | 2.081 | 26.489 | 0.047 | 0.037 |
| K09479 | ACADVL | Fire ~ TSF ~ Num. fires | 0.031 | 0.014 | 2.250 | 14.873 | 0.040 | 0.024 |
| K09699 | DBT/bkdB | Fire ~ TSF ~ Num. fires | -0.029 | 0.008 | -3.509 | 13.657 | 0.004 | 0.000 |
| K09699 | DBT/bkdB | TSF | -0.035 | 0.013 | -2.782 | 19.867 | 0.012 | 0.005 |
| K11645 | fdaB | Fire ~ TSF ~ Num. fires | -0.046 | 0.017 | -2.757 | 28.000 | 0.010 | 0.006 |
| K11645 | fdaB | TSF | -0.041 | 0.018 | -2.243 | 28.000 | 0.033 | 0.025 |
| K12308 | bgaB/lacA | Num. fires | 0.119 | 0.060 | 1.978 | 28.000 | 0.058 | 0.048 |
| K12960 | mtaD | Fire | -0.107 | 0.050 | -2.152 | 6.496 | 0.071 | 0.031 |
| K12960 | mtaD | Fire ~ Num. fires | -0.127 | 0.054 | -2.367 | 13.989 | 0.033 | 0.018 |
| K12960 | mtaD | Fire ~ TSF | 0.028 | 0.008 | 3.323 | 7.250 | 0.012 | 0.001 |
| K12960 | mtaD | Num. fires | -0.184 | 0.070 | -2.617 | 23.711 | 0.015 | 0.009 |
| K22224 | acdB | Fire ~ TSF | -0.023 | 0.008 | -2.756 | 10.463 | 0.019 | 0.006 |
| K22224 | acdB | TSF ~ Num. fires | -0.024 | 0.011 | -2.217 | 22.846 | 0.037 | 0.027 |
| K22224 | acdB | TSF ~ Num. fires | -0.024 | 0.011 | -2.217 | 22.846 | 0.037 | 0.027 |

| **Active taxa** |  |  |  |  |  |  |  |  |
| --- | --- | --- | --- | --- | --- | --- | --- | --- |
| **Phylum** | **Kingdom** | **Model term** | **LMM estimate** | **SE** | **t-statistic** | **df** | **P value (LMM)** | **P value (ANOVA - Type III)** |
| Basidiomycota | Fungi | Fire | -0.458 | 0.130 | -3.517 | 16.834 | 0.003 | 0.000 |
| Mucoromycota | Fungi | Fire | -0.307 | 0.089 | -3.463 | 28.000 | 0.002 | 0.001 |
| Chytridiomycota | Fungi | Fire | -0.188 | 0.095 | -1.982 | 28.000 | 0.057 | 0.048 |
| Verrucomicrobia | Bacteria | Fire | -0.091 | 0.046 | -1.965 | 9.687 | 0.079 | 0.049 |
| C. Bipolaricaulota | Bacteria | Fire | 0.081 | 0.039 | 2.055 | 19.951 | 0.053 | 0.040 |
| Firmicutes | Bacteria | Fire | 0.086 | 0.037 | 2.313 | 15.020 | 0.035 | 0.021 |
| Deinococcus Thermus | Bacteria | Fire | 0.098 | 0.045 | 2.208 | 10.093 | 0.052 | 0.027 |
| Bacteroidetes | Bacteria | Fire | 0.118 | 0.060 | 1.977 | 13.496 | 0.069 | 0.048 |
| C. Rokubacteria | Bacteria | Fire | 0.121 | 0.063 | 1.900 | 28.000 | 0.068 | 0.057 |
| C. division WS5 | Bacteria | Fire | 0.132 | 0.036 | 3.693 | 8.478 | 0.006 | 0.000 |
| Actinobacteria | Bacteria | Fire | 0.154 | 0.038 | 4.114 | 12.679 | 0.001 | 0.000 |
| C. Uhrbacteria | Bacteria | Fire | 0.171 | 0.084 | 2.037 | 28.000 | 0.051 | 0.042 |
| C. Parcubacteria | Bacteria | Fire | 0.183 | 0.043 | 4.239 | 11.474 | 0.001 | 0.000 |
| C. Marinimicrobia | Bacteria | Fire | 0.202 | 0.073 | 2.756 | 14.101 | 0.015 | 0.006 |
| Thaumarchaeota | Archaea | Fire | 0.228 | 0.097 | 2.339 | 28.000 | 0.027 | 0.019 |
| Crenarchaeota | Archaea | Fire | 0.234 | 0.066 | 3.574 | 28.000 | 0.001 | 0.000 |
| C. Peregrinibacteria | Bacteria | Fire | 0.269 | 0.070 | 3.841 | 10.780 | 0.003 | 0.000 |
| Cyanobacteria | Bacteria | Fire ~ Num. fires | -0.342 | 0.112 | -3.037 | 28.000 | 0.005 | 0.002 |
| Deinococcus Thermus | Bacteria | Fire ~ Num. fires | -0.142 | 0.045 | -3.154 | 19.636 | 0.005 | 0.002 |
| Fusobacteria | Bacteria | Fire ~ Num. fires | -0.075 | 0.028 | -2.650 | 25.131 | 0.014 | 0.008 |
| Crenarchaeota | Archaea | Fire ~ Num. fires | 0.132 | 0.059 | 2.237 | 28.000 | 0.033 | 0.025 |
| C. Bipolaricaulota | Bacteria | Fire ~ Num. fires | 0.135 | 0.035 | 3.806 | 25.999 | 0.001 | 0.000 |
| C. Uhrbacteria | Bacteria | Fire ~ Num. fires | 0.188 | 0.075 | 2.494 | 28.000 | 0.019 | 0.013 |
| Chloroflexi | Bacteria | Fire ~ Num. fires | 0.246 | 0.062 | 3.983 | 23.244 | 0.001 | 0.000 |
| Fusobacteria | Bacteria | Fire ~ TSF | -0.013 | 0.005 | -2.768 | 19.289 | 0.012 | 0.006 |
| C. Woesearchaeota | Archaea | Fire ~ TSF | 0.016 | 0.005 | 3.375 | 11.736 | 0.006 | 0.001 |
| Thermotogae | Bacteria | Fire ~ TSF | 0.025 | 0.012 | 2.097 | 18.942 | 0.050 | 0.036 |
| C. division WWE3 | Bacteria | Fire ~ TSF | 0.034 | 0.014 | 2.459 | 28.000 | 0.020 | 0.014 |
| Verrucomicrobia | Bacteria | Fire ~ TSF ~ Num. fires | -0.025 | 0.008 | -3.138 | 16.582 | 0.006 | 0.002 |
| C. Peregrinibacteria | Bacteria | Fire ~ TSF ~ Num. fires | -0.024 | 0.011 | -2.124 | 18.395 | 0.047 | 0.034 |
| Cryptomycota | Fungi | Fire ~ TSF ~ Num. fires | 0.018 | 0.007 | 2.721 | 15.115 | 0.016 | 0.007 |
| Cyanobacteria | Bacteria | Num. fires | -0.313 | 0.112 | -2.780 | 28.000 | 0.010 | 0.005 |
| Basidiomycota | Fungi | Num. fires | -0.279 | 0.131 | -2.139 | 26.119 | 0.042 | 0.032 |
| Verrucomicrobia | Bacteria | Num. fires | -0.152 | 0.073 | -2.085 | 25.813 | 0.047 | 0.037 |
| C. Woesearchaeota | Archaea | Num. fires | 0.075 | 0.038 | 1.972 | 25.725 | 0.059 | 0.049 |
| Proteobacteria | Bacteria | Num. fires | 0.111 | 0.039 | 2.839 | 25.430 | 0.009 | 0.005 |
| C. Kaiserbacteria | Bacteria | Num. fires | 0.120 | 0.062 | 1.948 | 24.961 | 0.063 | 0.051 |
| Deferribacteres | Bacteria | Num. fires | 0.148 | 0.060 | 2.456 | 28.000 | 0.021 | 0.014 |
| C. Bipolaricaulota | Bacteria | Num. fires | 0.148 | 0.036 | 4.153 | 26.117 | 0.000 | 0.000 |
| Crenarchaeota | Archaea | Num. fires | 0.156 | 0.059 | 2.644 | 28.000 | 0.013 | 0.008 |
| Chloroflexi | Bacteria | Num. fires | 0.227 | 0.070 | 3.260 | 26.144 | 0.003 | 0.001 |
| Verrucomicrobia | Bacteria | TSF | -0.043 | 0.014 | -2.992 | 22.609 | 0.007 | 0.003 |
| Cyanobacteria | Bacteria | TSF | -0.040 | 0.020 | -2.012 | 28.000 | 0.054 | 0.044 |
| C. Portnoybacteria | Bacteria | TSF | -0.022 | 0.008 | -2.695 | 28.000 | 0.012 | 0.007 |
| C. division Kazan 3B28 | Bacteria | TSF | 0.021 | 0.009 | 2.329 | 28.000 | 0.027 | 0.020 |
| Chloroflexi | Bacteria | TSF | 0.038 | 0.013 | 2.962 | 21.935 | 0.007 | 0.003 |
| Euryarchaeota | Archaea | TSF ~ Num. fires | -0.034 | 0.016 | -2.169 | 24.682 | 0.040 | 0.030 |
| C. Nealsonbacteria | Bacteria | TSF ~ Num. fires | -0.031 | 0.016 | -1.937 | 24.165 | 0.065 | 0.053 |
| C. Portnoybacteria | Bacteria | TSF ~ Num. fires | -0.016 | 0.007 | -2.250 | 28.000 | 0.033 | 0.024 |
| C. Parcubacteria | Bacteria | TSF ~ Num. fires | 0.020 | 0.008 | 2.464 | 24.362 | 0.021 | 0.014 |
| C. Uhrbacteria | Bacteria | TSF ~ Num. fires | 0.029 | 0.012 | 2.491 | 28.000 | 0.019 | 0.013 |

**Supplemental Table 4**. Summary of KEGG functional pathway enrichment analysis within modules of control (no fire), post-fire (fire), and time since fire (TSF) weighted gene co-expression networks.

| **Module** | **Enriched KEGG Pathways** | **Total KOs** | **Shared KOs** | **Unique KOs** | **Shared KOs (%)** |
| --- | --- | --- | --- | --- | --- |
| Unburned | 62 | 1046 | 545 | 501 | 52.1 |
| Burned | 25 | 189 | 81 | 108 | 42.8 |
| TSF | 4 | 29 | 7 | 22 | 24.1 |

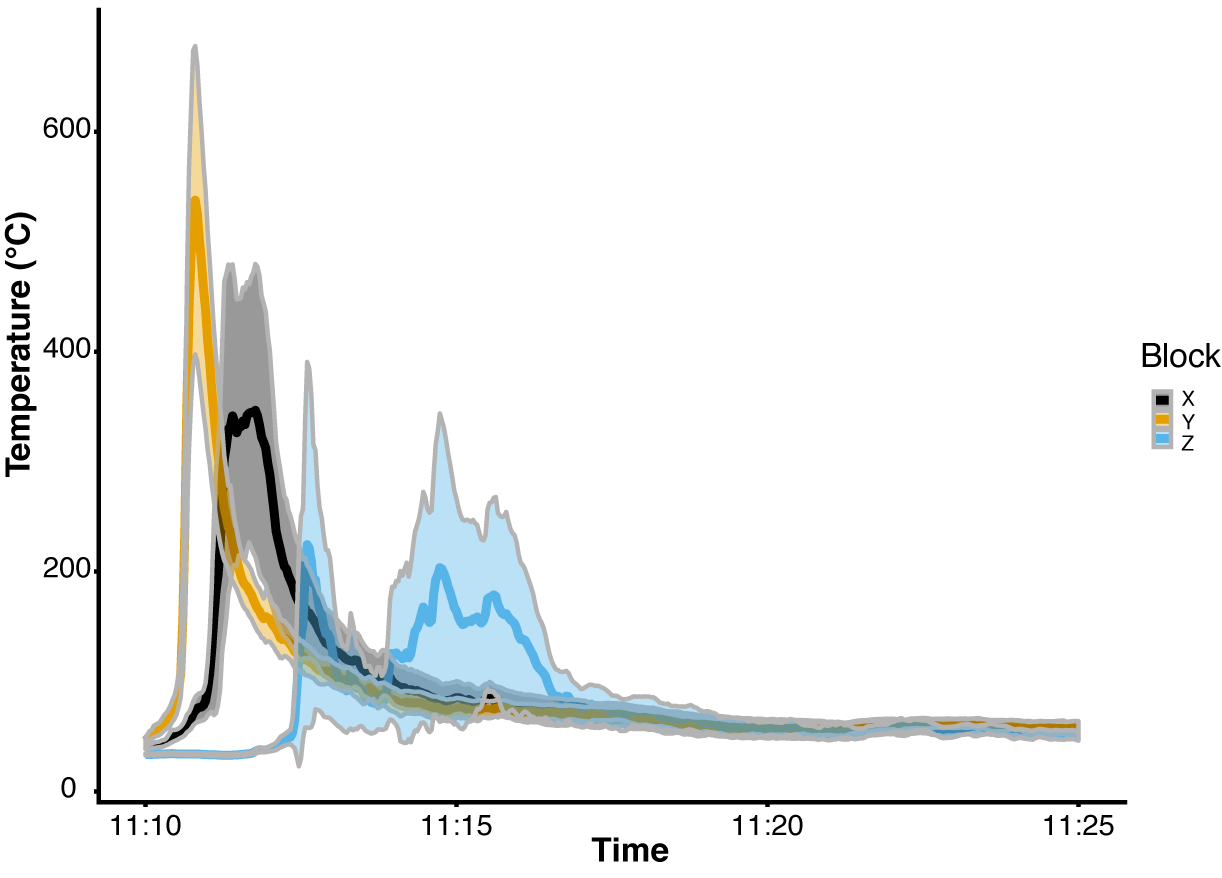

**Supplemental Figure 1**. Soil surface temperatures experienced during prescribed burn treatment over time. Samples analyzed for this study were chosen from block Y (yellow), which across the sampled locations maintained the most consistently high temperatures.

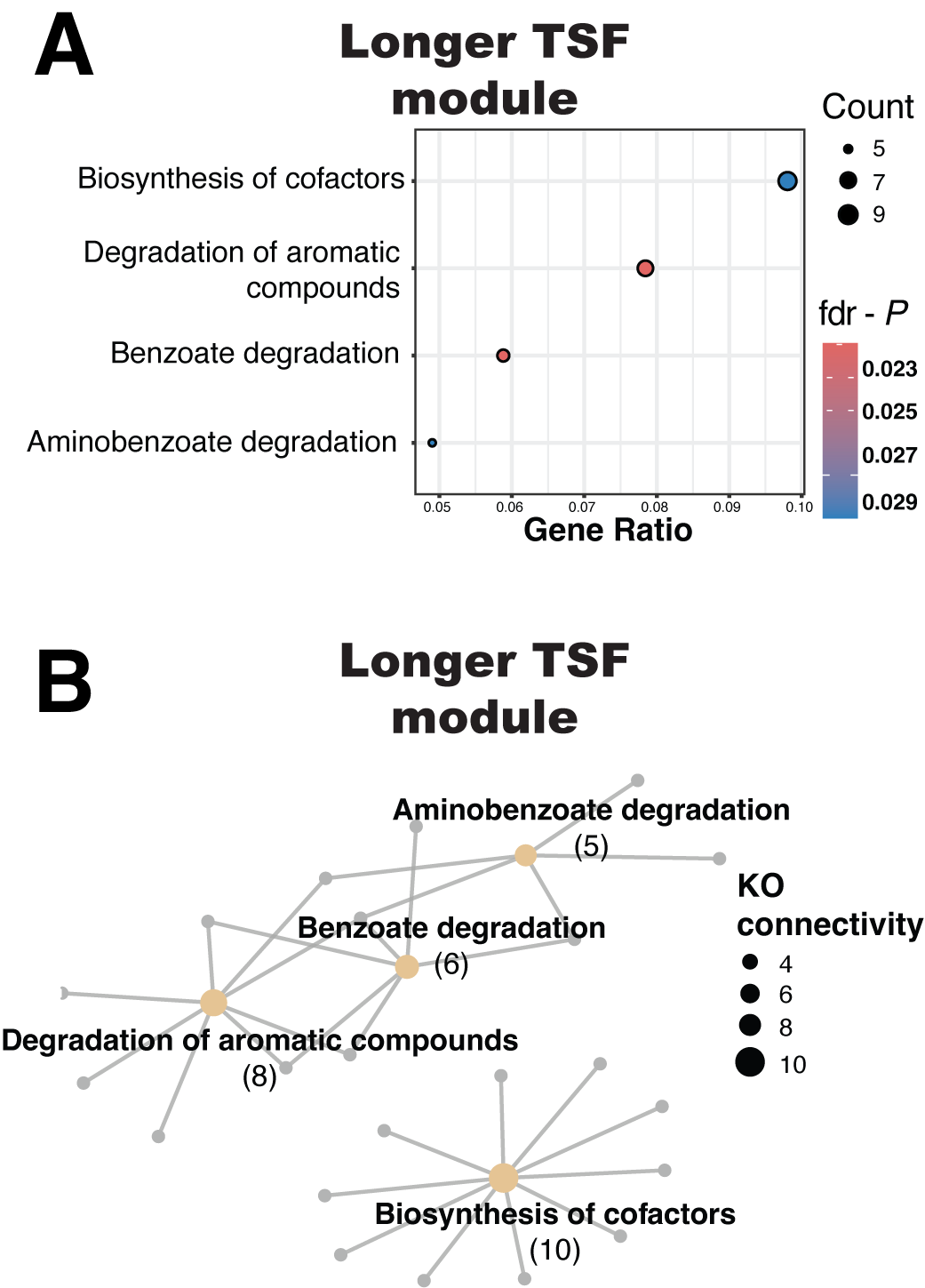

**Supplemental Figure 2**. Functional pathways enriched in the weighted gene co-expression network module associated with time since fire (TSF). Gene ratios are plotted for the four representative pathways identified in the module (A). Network diagrams show functional pathways in yellow and associated KEGG orthologs (genes) in grey (B).

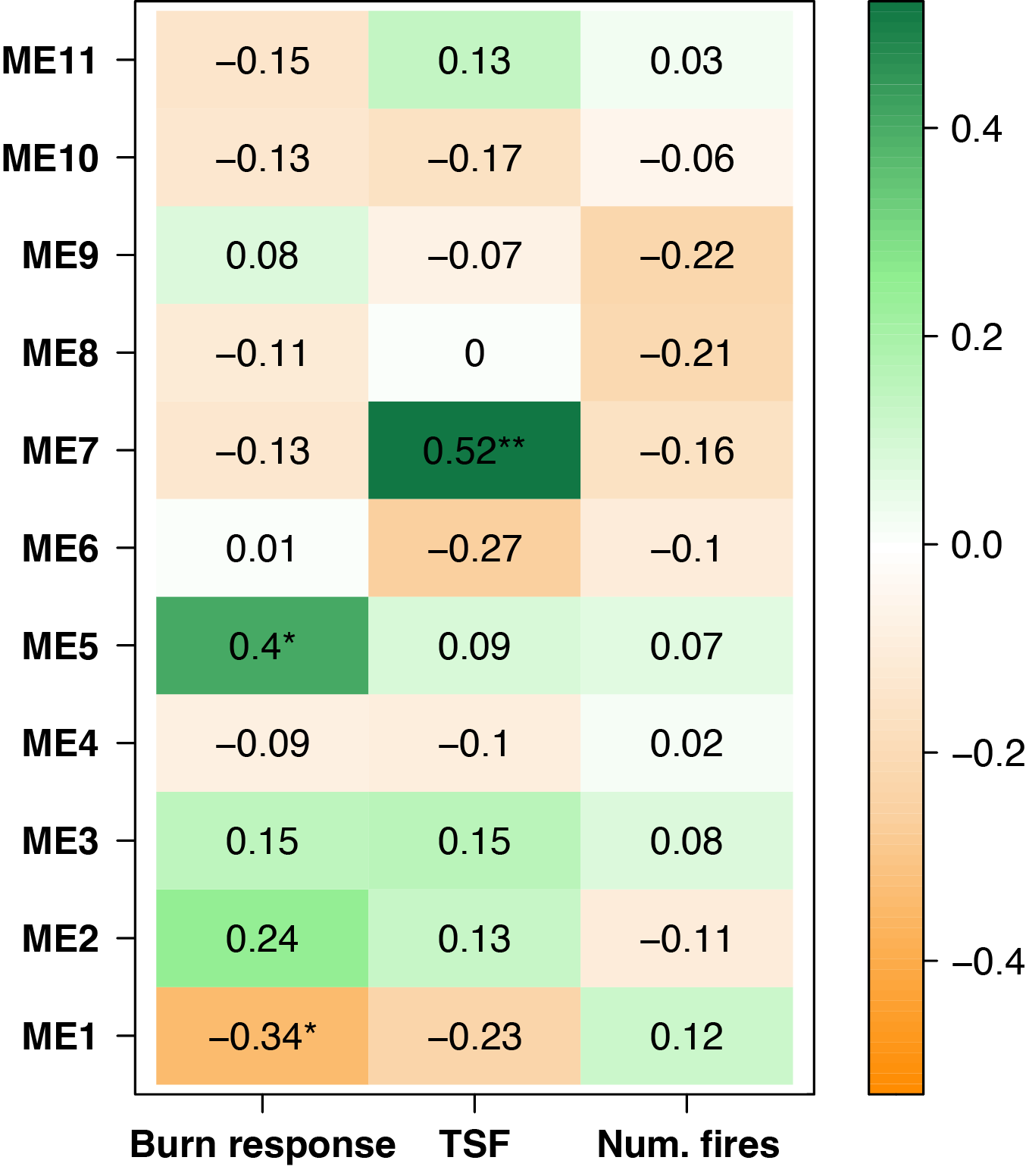

**Supplemental Figure 3**. Correlation coefficients (Spearman’s rho) between experimental treatments: prescribed fire (Burn response), time since fire (TSF), historic number of fires (Num. fires), and weighted gene co-expression network module eigengene (ME). Burn response of one is equal to burning, while burn response of zero is equal to unburned, control soils. * = *P* < 0.05, ** = *P* < 0.01. ME1 and ME5 were representative of the ‘unburned’ and ‘burned’ treatments, respectively, while ME7 responded positively to increasing TSF.

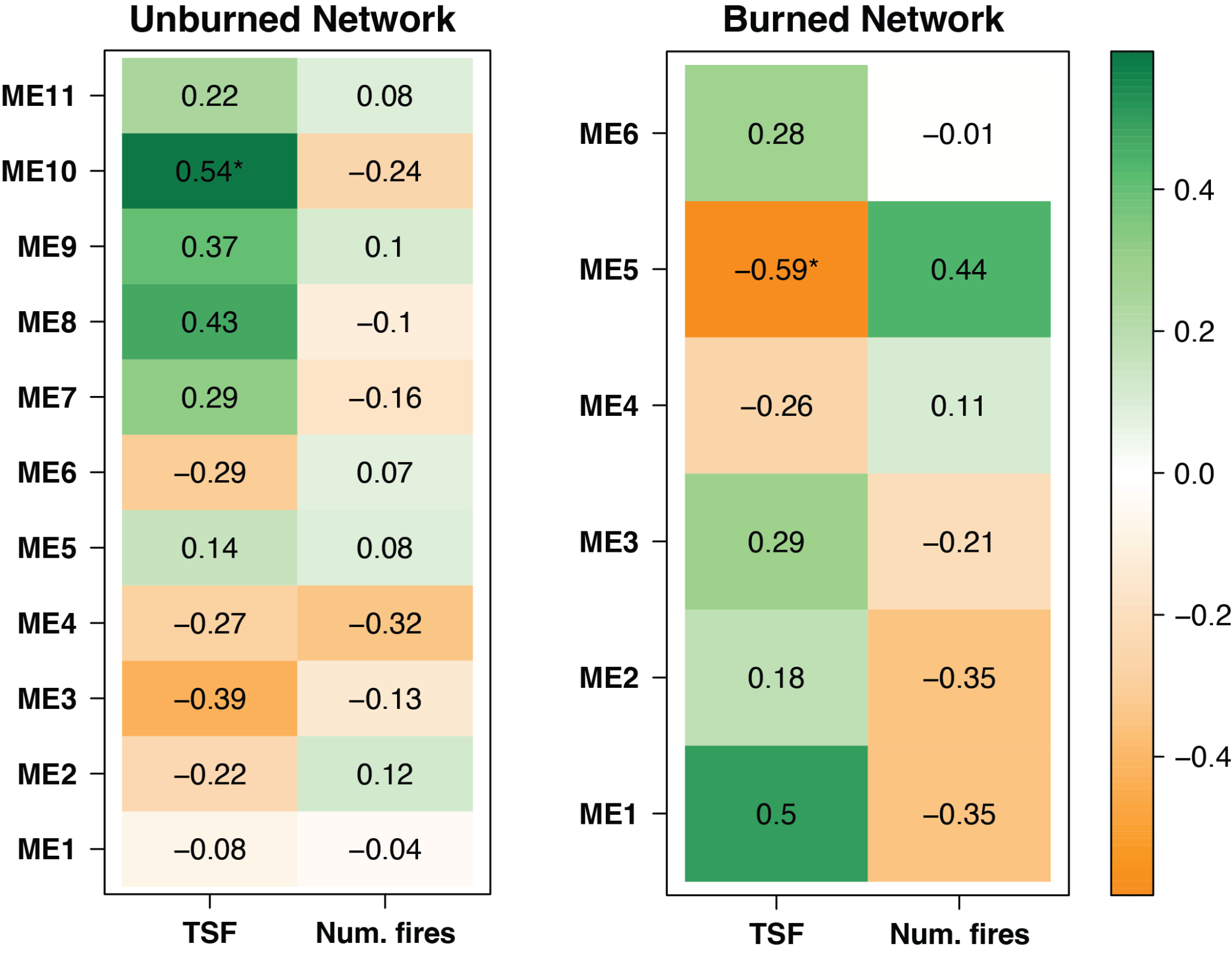

**Supplemental Figure 4**. Module eigengenes (ME) identified within either unburned or burned weighted gene co-expression networks and their association with fire legacy metrics: time since fire (TSF) and historic number of fires (Num. fires). Color represents Spearman’s rho correlation coefficient, and * = *P* < 0.05. This analysis highlights the simplification of gene co-expression after fire (halving the number of module eigengenes).

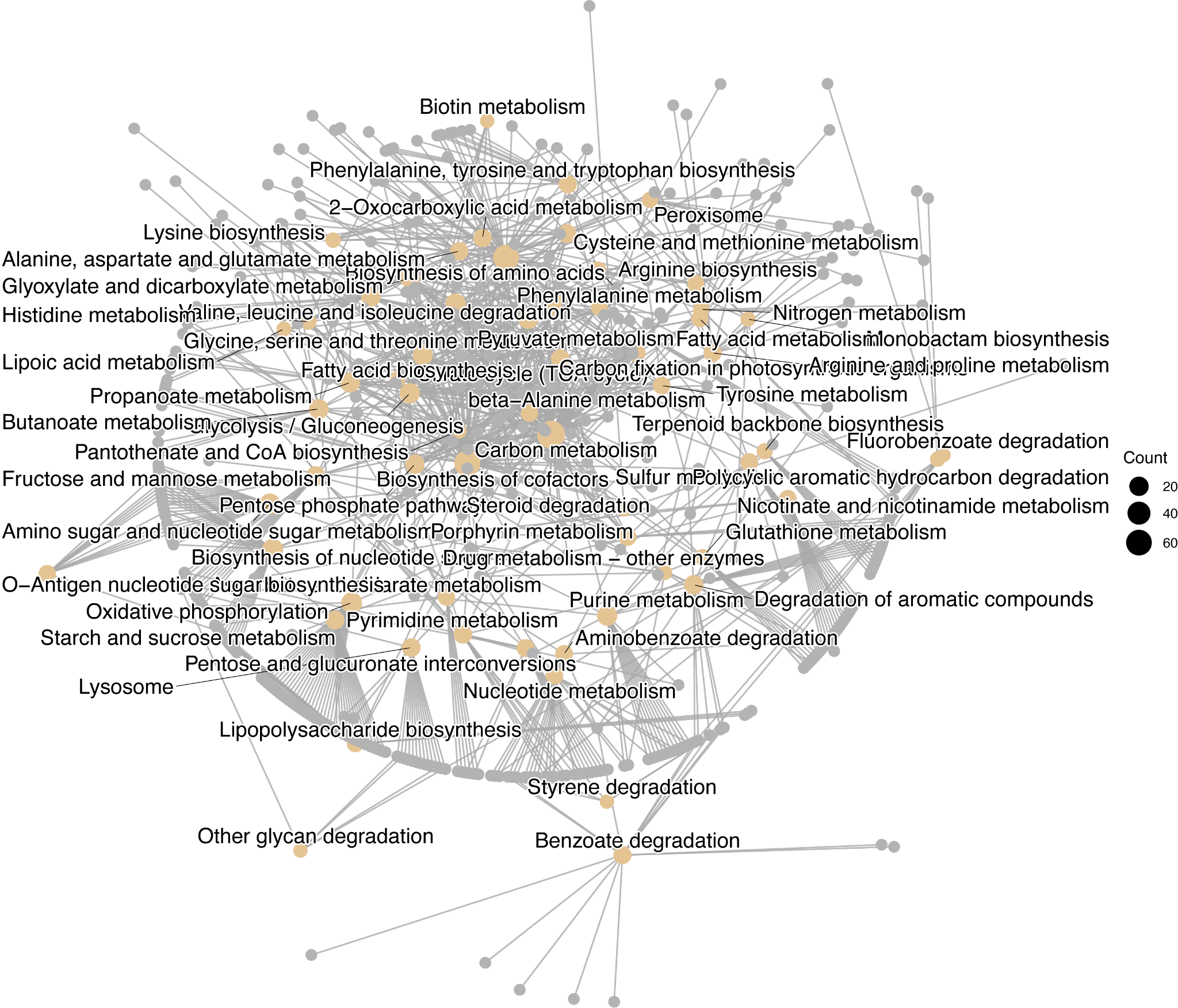

**Supplemental Figure 5**. Functional pathways enriched in the weighted gene co-expression network module associated with unburned, control soil metatranscriptomes. All 62 representative pathways are plotted in the network diagram, showing functional pathways in yellow and associated KEGG orthologs (genes) in grey.

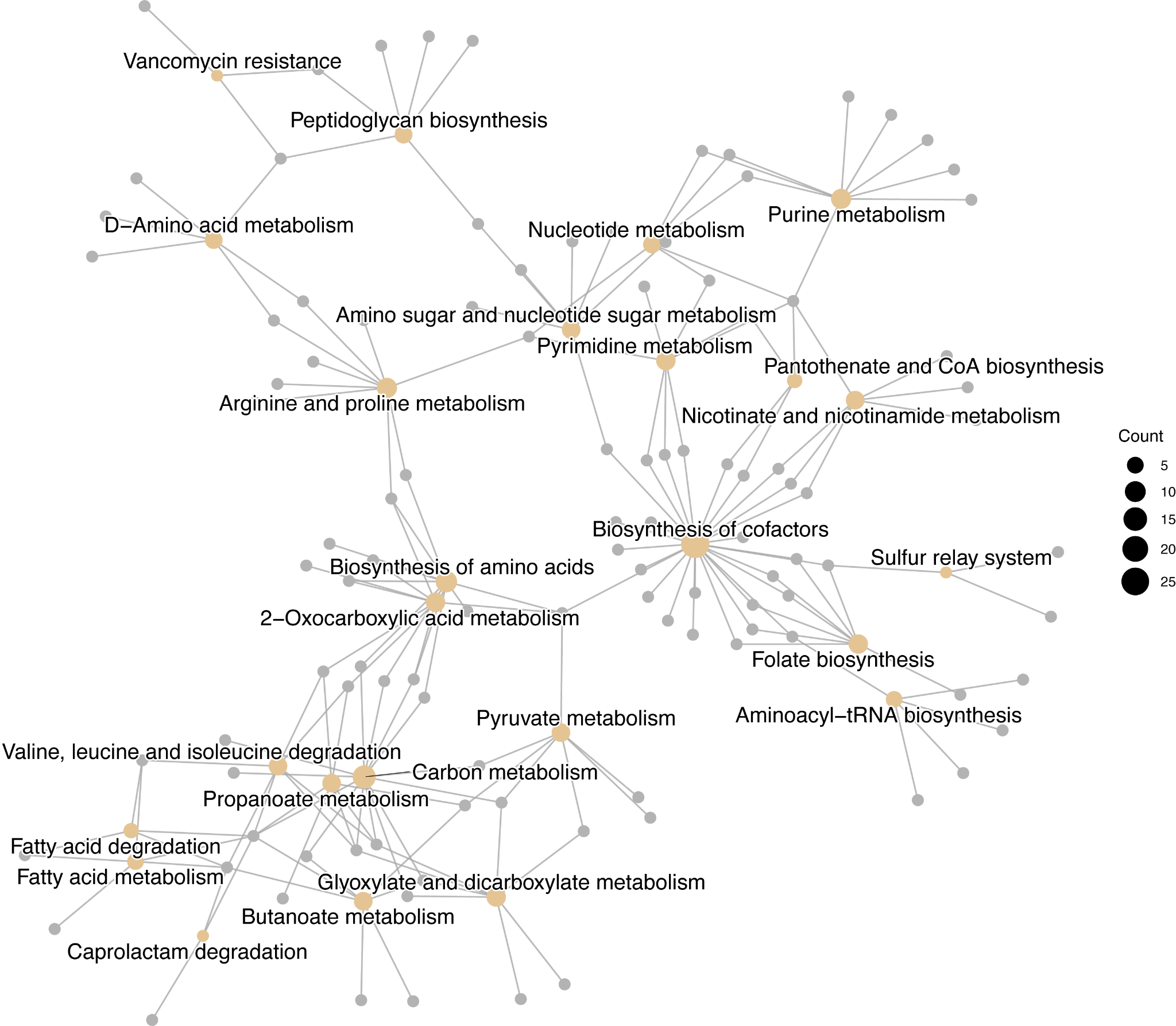

**Supplemental Figure 6**. Functional pathways enriched in the weighted gene co-expression network module associated with post-fire soil metatranscriptomes. All 25 representative pathways are plotted in the network diagram, showing functional pathways in yellow and associated KEGG orthologs (genes) in grey.

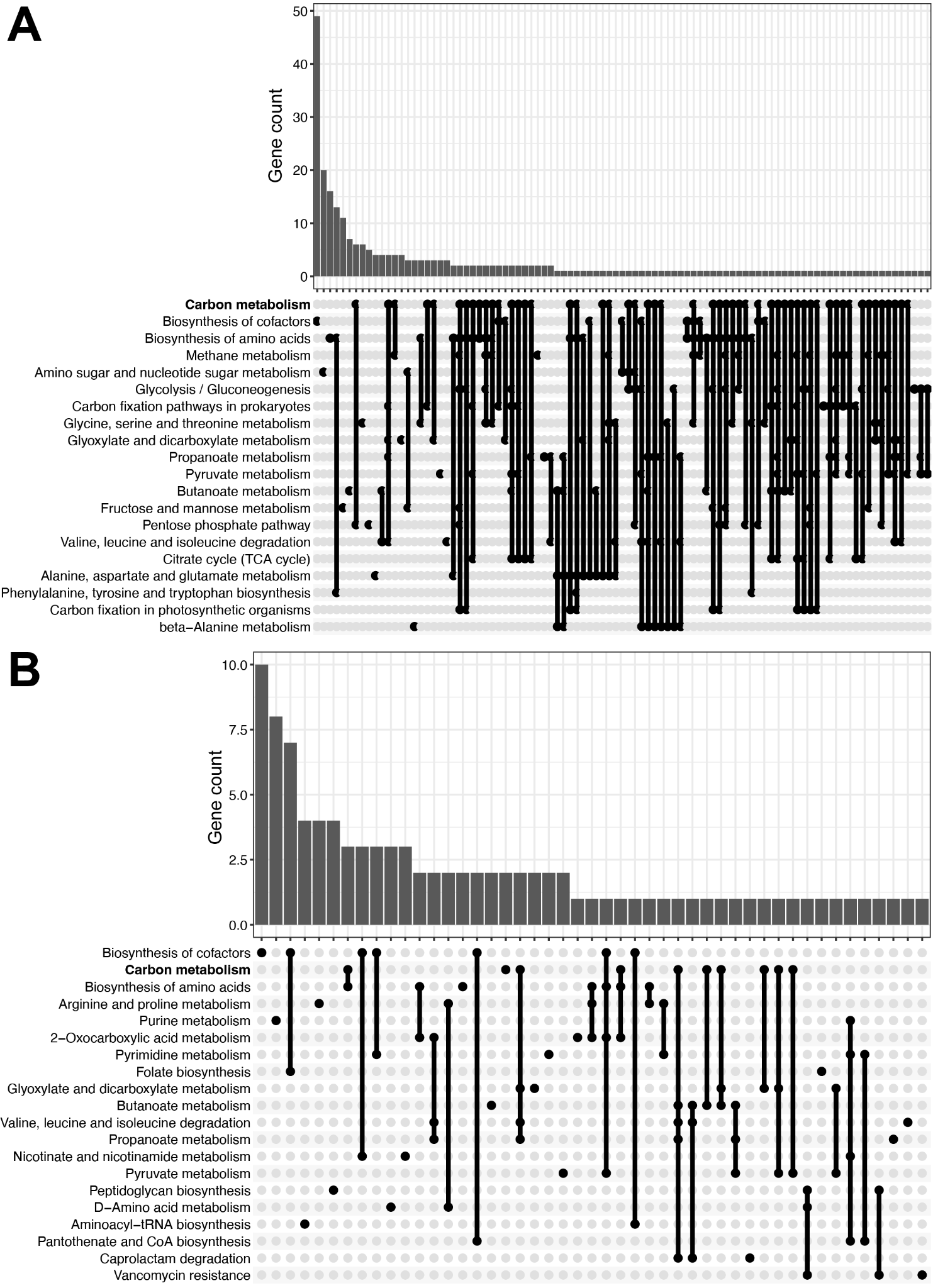

**Supplemental Figure 7**. Functional pathways enriched in weighted gene co-expression network modules associated with unburned or experimentally burned soils. Upset plots for the 20 largest pathways (by gene count) in unburned control module (A) and burned soil module (B). Functional pathways are organized on the y axis by KEGG gene count (top-to-bottom). Note differences in KEGG ortholog count within functional pathways for A and B. This plot highlights the dramatic loss of metatranscriptome co-expression network functional pathway interconnectivity after fire. Carbon metabolism in bold was the most significantly enriched functional pathway in unburned control soils, while it was not significantly enriched after burning (despite a relatively high gene count).
